# Informational Masking Constrains Vocal Communication in Nonhuman Animals

**DOI:** 10.1101/2022.03.31.486641

**Authors:** Saumya Gupta, Lata Kalra, Luke Drummer, Gary J. Rose, Mark A. Bee

**Affiliations:** Department of Ecology, Evolution, and Behavior, University of Minnesota, Saint Paul, MN 55108, USA; School of Biological Sciences, University of Utah, Salt Lake City, UT 84112, USA; Graduate Program in Neuroscience, University of Minnesota, Minneapolis, MN 55455, USA

## Abstract

Animals often communicate acoustically in noisy social environments, yet how receivers extract relevant information from overlapping sounds is poorly understood. Studies of animal communication in noise are typically based on a traditional filterbank model of hearing focused on *energetic masking*, where spectrotemporal overlap in peripheral auditory filters limits signal audibility. By contrast, studies of human speech show that background sounds can also interfere with selecting and attending to otherwise audible signals in a phenomenon known as *informational masking*. Whether informational masking constrains acoustic communication in nonhuman animals remains poorly understood. Through controlled laboratory experiments with treefrogs (*Hyla chrysoscelis*), we demonstrate, for the first time in a nonhuman animal, that informational masking can impair crucial mate-choice decisions when vocal signals and other concurrent sounds share similar temporal features. These impacts spanned a wide range of signal-to-noise ratios, occurred in the absence of spectrotemporal overlap in the auditory periphery, and were strongest when interfering sounds fell within a frequency band salient for vocal processing. Our findings challenge conventional views on how noise impacts animal communication by establishing informational masking as a general communication problem shared by humans and other animals. These results highlight the need to distinguish between energetic and informational masking to understand the evolution of signaling in complex acoustic environments.

## Introduction

Noise is a potent source of selection on animal communication (*1–3*). By inducing costly communication errors, auditory masking negatively impacts consequential behaviors with direct and profound influences on survival and reproduction, such as mate choice (*4, 5*), prey detection (*6, 7*), and threat evasion (*8*). Not surprisingly, animals exhibit a diversity of adaptations for coping with noise problems. These adaptations include, for example, shifting the spectral frequencies of signals away from those of dominant noise sources (*9, 10*), signaling at higher amplitudes in noise (*11, 12*), and signaling in locations or at times when interfering noise sources are less prevalent (*13–15*).

Our current understanding of how animals are adapted to cope with noise is largely informed by Fletcher’s influential filterbank model of the auditory system (*16*). Applied across taxa, the model conceptualizes peripheral auditory systems as one or more bandpass filters, each tuned to a specific frequency range, that function to decompose received sounds into their spectral components for further processing by the central nervous system. In this model, *energetic masking* occurs when a signal’s audibility is reduced because competing sounds that are transduced by the same auditory filter as the signal and overlap with it in time add enough energy to raise the threshold signal-to-noise ratio (SNR) for detection (*17, 18*). Across diverse animal taxa, the adaptations for communicating in noise identified thus far share a common function: to reduce energetic masking by improving the SNR for communication at the auditory periphery.

The explicit or implicit focus on peripheral filtering and energetic masking that underlies most studies of animal communication overlooks a critical dimension of the noise problem: competing sounds can impair signal perception even when they are spectrally distinct and thus separable in the auditory periphery (*19, 20*). As identified by studies of human hearing and speech communication, this second and fundamentally different process, known as *informational masking*, occurs when central auditory processes fail to correctly extract information from an otherwise audible signal (*19, 20*). Informational masking is thought to arise from constraints on auditory stream segregation or selective attention, and is often most pronounced when signals and competing sounds share acoustic features in common, such as similar temporal patterns (*21*), a common spatial origin (*22*), or similar linguistic content in the case of speech (*23*). Unlike energetic masking, informational masking does not always decrease with increasing SNR and can persist even at high SNRs, making its effects potentially more severe and less predictable (*24, 25*). Importantly, informational masking is recognized as a major contributor to the human “cocktail party problem,” which highlights the difficulty of understanding speech when there are multiple competing speakers (*24, 26, 27*).

The distinction between energetic and informational masking forces a critical re-evaluation of noise-driven selection on animal communication systems. Compared with our current understanding of how informational masking impacts speech perception, we know far less about how informational masking might impact communication in other animals (*28, 29*). This gap in knowledge limits our ability to elucidate the mechanisms and evolution of acoustic communication across diverse animal species, particularly those that communicate in “cocktail-party-like” listening environments, such as insect and frog breeding choruses (*30, 31*), songbird dawn choruses (*32*), penguin crèches (*33*), and groups of foraging bats (*34*). In such species, the costs of informational masking might be particularly high, because receivers frequently encounter situations where multiple individuals in close proximity simultaneously produce concurrent signals with similar acoustic features (*35*). If informational masking is a widespread phenomenon, it could represent a potent selective pressure shaping animal communication systems that has thus far been largely ignored.

Here, we use biologically relevant target signals and concurrent sounds to test the hypothesis that informational masking can impair vocal communication in a nonhuman animal. Across species, temporal modulations of the amplitude envelopes of communication signals convey key biological information (*36*), and they are critical for human speech perception (*37*) as well as signal recognition and discrimination in a diversity of nonhuman animals (*38, 39*). The taxonomically widespread use of temporal patterns in communication signals suggests central mechanisms for temporal processing could be particularly susceptible to informational masking in social environments, such as frog breeding choruses, where many individuals produce concurrent signals with broadly similar temporal features. Using female phonotaxis as a behavioral assay, we investigated informational masking in Cope’s gray treefrog, *Hyla chrysoscelis*, a well-studied anuran species in which females rely on temporal features of male vocal signals to choose a mate in the noisy environment of a breeding chorus (*30, 38, 40*).

## Results

### Does Informational Masking Reduce Sound-evoked Mate Choice Behavior**?**

In Cope’s gray treefrogs, males form dense, mixed-species choruses where they produce loud advertisement calls (Fig. 1A) with a species-specific temporal modulation structure that is key to attracting conspecific females. Each call consists of a train of 12 to 43 evenly spaced pulses, with an average pulse rate of about 50 pulses/s (*41*). High levels of concurrent calling by conspecific males, as well as males of different species that also produce pulsatile calls within the gray treefrog hearing range, are commonplace in treefrog breeding choruses (*42*). Mating commences when a female exhibits a stereotyped and well-characterized behavior known as phonotaxis (approach toward sound) in response to hearing the advertisement calls of a stationary conspecific male in a chorus (*43*). Thus, in choosing a mate, females must correctly exhibit phonotaxis behavior in response to the temporally modulated calls of a conspecific in a listening environment awash with concurrent sounds with similar temporal features. Based on this natural acoustic context, we evaluated whether concurrent sounds with similar temporal patterns could disrupt mate choice behavior in a way that would be expected if treefrogs were susceptible to informational masking.

**Fig. 1.**
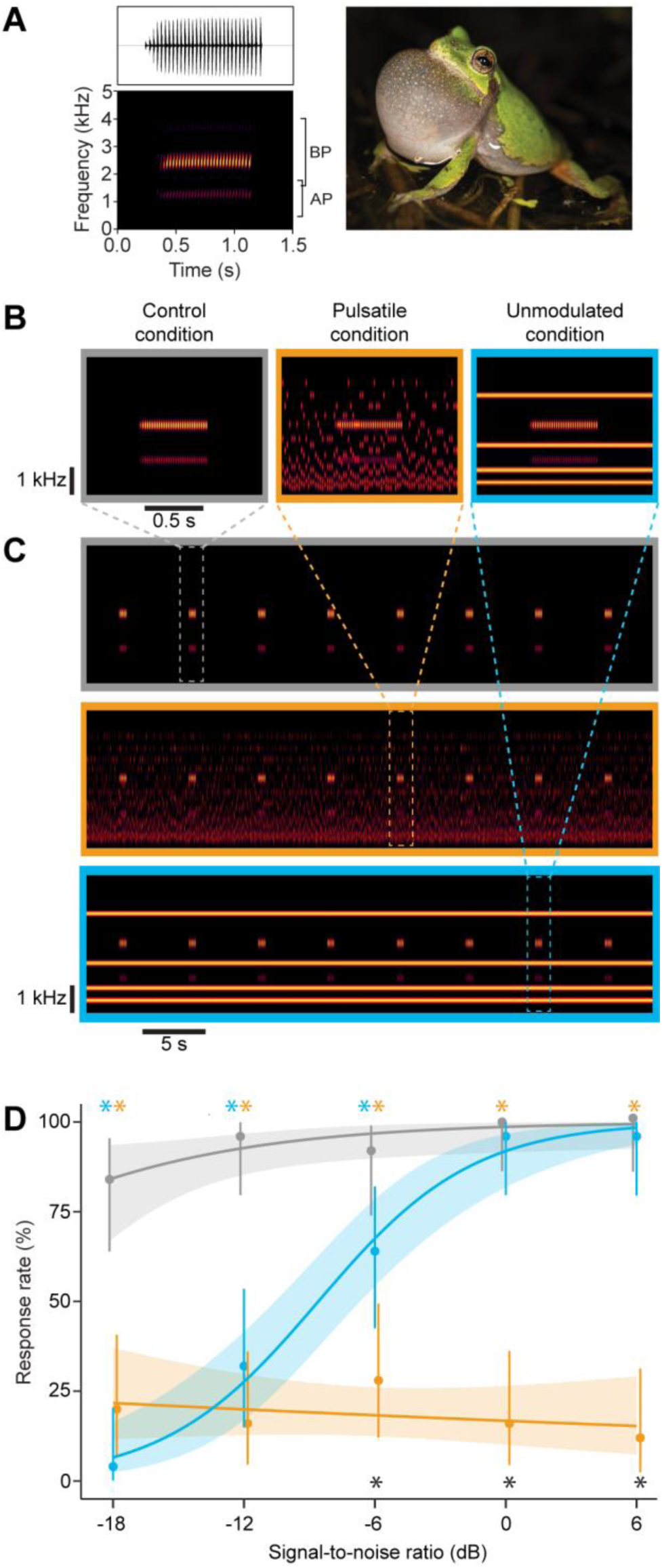
Experiment 1: Gray treefrogs experience greater auditory masking when concurrent sounds are temporally modulated. (**A**) Spectrogram (bottom) and waveform (top) illustrating the pulsatile temporal envelope and frequency composition of a natural advertisement call of *Hyla chrysoscelis* and a picture of a calling male (photo credit: Alex Baugh). (**B**) Spectrograms illustrating the synthetic call used as a target signal in Experiment 1 in the control (*gray*), pulsatile (*orange*), and unmodulated (*blue*) conditions. In Experiment 1, the pulsatile and unmodulated concurrent sounds were broadcast at 79 dB SPL; signal amplitude was varied to achieve a signal-to-noise ratio (SNR) from -18 to +6 dB. (**C**) Three 45-s segments of stimuli used during the no-choice tests of Experiment 1, in which the target signal was repeated at a rate of 11 calls/min to simulate a naturally calling male in the control (*gray*), pulsatile (*orange*), and unmodulated (*blue*) conditions. (**D**) Response rates showing the percentages (± 95% exact binomial confidence intervals) of subjects (*n* = 25 per condition) that responded to the target signal in the control, pulsatile, and unmodulated conditions as a function of SNR. Points indicate observed percentages. Solid lines represent 95% exact binomial confidence intervals. Shaded regions show predicted response probabilities from generalized linear model. Female response rates were uniformly high when target signals were presented in the control condition. In the unmodulated condition, response rates were low at the lowest SNR and increased monotonically as SNR increased. In the pulsatile condition, response rates were low and did not change with increasing SNR. Blue and orange asterisks depict significant differences between the control and the unmodulated and pulsatile conditions, respectively; black asterisks indicate significant differences between the unmodulated and pulsatile conditions (see Table S1 for details).

#### Experiment 1: Impacts of Concurrent Sounds with Temporal Modulations on Behavioral Responses to Vocal Signals

In Experiment 1, we tested the hypothesis that temporal modulations in concurrent sounds – a known cause of informational masking in humans (*21*) – reduce acoustically-evoked phonotaxis behavior beyond what can be explained by energetic masking alone. In a behavioral test that presented sounds from a single speaker (Fig. S1A), we measured female phonotaxis in response to a simulated male calling in a control condition (signal-alone) and in the presence of two different concurrent sounds. The target signal (Fig. 1B) was a synthesized advertisement call based on the average properties of calls recorded in local populations (*41*). The call repeated at a rate of 11 calls/min and consisted of a sequence of 30 identical pulses delivered at 50 pulses/s, each having two harmonically related spectral components with frequencies (and relative amplitudes) of 1.25 kHz (-11 dB) and 2.5 kHz (0 dB). Each concurrent sound played continually during an experimental trial to simulate the continuous sounds present in choruses (Fig. 1B, 1C). Both concurrent sounds consisted of a 4-component harmonic complex (harmonics 1, 2, 4, and 8) spanning the gray treefrog hearing range (*44*). The amplitude envelope of the concurrent sounds either lacked temporal modulations (the unmodulated condition) or consisted of a series of pulses that had the same temporal structure as the pulses in the target signal, with which they were interleaved in time (the pulsatile condition) (Fig. 1B, 1C). A key feature of this stimulus design is that the unmodulated concurrent sound overlapped in time with pulses in the target signal, whereas the pulses in the pulsatile concurrent sound were temporally interleaved with those of the signal, thereby avoiding simultaneous overlap and minimizing energetic masking. Each concurrent sound was broadcast at 79 dB SPL (LC_eq_), and the amplitude of the target signal was varied across trials to produce SNRs (or equivalent amplitudes in the control condition) ranging from -18 dB to +6 dB. When presented by themselves in additional sham conditions (i.e., no-signal), neither concurrent sound evoked positive or negative phonotaxis (Fig. S2A). We predicted phonotaxis behavior would be reduced in the pulsatile condition relative to the unmodulated condition if temporal modulations in concurrent sounds contributed to informational masking.

Our results revealed a clear divergence in phonotaxis responses between acoustic conditions. In the control condition, females exhibited robust phonotaxis, with response rates increasing from 84% to 100% with increasing signal amplitude (GEE: β = 0.006, SE = 0.002, *P* = 0.0150; Fig. 1D). In the unmodulated condition, response rates increased steeply with SNR from 4% at -18 dB to 96% at +6 dB (GEE: β = 0.041, SE = 0.002, *P* < 0.0001; Fig.1D). Compared with the control condition, response rates were significantly lower in the unmodulated conditions at lower SNRs (≤ -6 dB) but not at higher SNRs (≥ 0 dB) (Fig. 1D, Table S1). In stark contrast, phonotaxis response rates in the pulsatile condition remained consistently low (between 12-28%) across the entire SNR range and showed no significant recovery as signal amplitude increased (GEE: β = -0.003, SE = 0.004, *P* = 0.5130; Fig.1D). Response rates were significantly lower at all SNRs in the pulsatile condition compared to control condition (Fig. 1D, Table S1). Compared with the unmodulated condition, response rates in the pulsatile condition were similar at low SNRs (≤ -12 dB) and significantly lower at higher SNRs (≥ -6 dB) (Fig. 1D, Table S1).

Overall, these results suggest subjects experienced informational masking in the presence of concurrent sounds that were temporally modulated. Consistent with earlier studies of masking in gray treefrogs (*30*), both the unmodulated and pulsatile concurrent sounds reduced responses to target signals at low SNRs. However, subjects experienced a release from masking as SNR increased in the unmodulated condition, but not the pulsatile condition. This observed pattern of performance improving with increasing SNR in the unmodulated condition, but remaining persistently poor even at higher SNR in the pulsatile condition, parallels known differences in the effects of energetic and informational masking, respectively, in human studies (*24, 25*). However, results from Experiment 1 do not conclusively rule out energetic masking because the signal and pulsatile concurrent sounds had overlapping frequency spectra and identical modulation rates (Figure 1B). Thus, reduced phonotaxis in the pulsatile condition might have resulted from either energetic masking in the modulation domain (*45*) or to non-simultaneous energetic masking (*46*), whereby the interleaved pulses of the concurrent sound elicited forward masking of pulses in the signal despite not overlapping in time. To eliminate these possibilities, and to further investigate the potential impacts of informational masking, we conducted two additional experiments. In these experiments, we minimized overlap between signals and concurrent sounds in both the frequency and modulation domains to assess how spectrally remote sounds modulated at different rates affect call recognition and discrimination.

### Does Informational Masking Disrupt Recognition and Discrimination of Vocal Signals?

Vocal signals mediate species recognition and sexual selection in gray treefrogs, in which females exhibit selective phonotaxis based on the temporal structure of vocalizations (*38, 40, 43, 47, 48*). In both Cope’s gray treefrog (*H. chrysoscelis*, a diploid) and in its tetraploid sister species (*H. versicolor*), females only recognize sounds as conspecific calls if they exceed a threshold number of pulses in duration (*48*). For calls surpassing this threshold, females discriminate against short calls in favor of longer calls with more pulses. In the tetraploid, the metabolic demands of calling increase with call duration (*49*), and males that produce longer calls sire offspring with higher fitness, suggesting pulse number is an indicator of male quality (*50*).

We tested the hypothesis that informational masking disrupts the ability of females to recognize and discriminate among conspecific signals based on pulse number. To do so, we used stimuli that exploited two key features of hearing and communication in gray treefrogs to minimize energetic masking. First, the two harmonically related spectral components in gray treefrog advertisement calls primarily stimulate two physically distinct sensory organs in the frog inner ear (*38, 44*) (Figs. 1A, S3). The amphibian papilla (AP) is tonotopically organized and tuned to lower frequencies, whereas the basilar papilla (BP) lacks tonotopy and is tuned to relatively higher frequencies. In Cope’s gray treefrogs, the best excitation frequencies of the AP and BP are approximately 1.4 kHz and 2.5 kHz, respectively, with minimal overlap in the tuning of the two papilla (*44*). The lower (∼1.25 kHz) spectral component of the call primarily stimulates the AP and the higher (∼2.5 kHz) component stimulates the BP. Second, although natural calls have both spectral components (Fig. 1A) (*41*), females do not require both components to recognize conspecific calls. Females readily respond to synthesized calls having species-specific pulse rates but just one of the two spectral components (Fig. S3A) (*47, 51, 52*). These two features allowed us to minimize energetic masking by experimentally placing target signals and concurrent sounds in different frequency ranges such that each was primarily transduced by a physically distinct inner ear organ.

In Experiments 2 and 3, the frequency of each pulse in the target signal (50 pulses/s) was always fixed at either 1.25 kHz (AP range) or 2.5 kHz (BP range), and concurrent sounds were restricted to the opposite frequency range (Fig. S3). We designed a “pulsatile condition” (Fig. S3B) that combined elements of informational maskers from classic experiments on humans (e.g., the multitone masking paradigm) (*53*) with biologically realistic features of frog choruses (*42*). In this condition, the concurrent sound was a gated series of pulses in which the frequency of each pulse varied randomly within a narrow range to stimulate the AP or BP; these stimuli were designed to partially simulate a concurrent call produced by a heterospecific frog calling nearby. Pulses were presented at 25 pulses/s (i.e., half the target signal’s pulse rate) and temporally interleaved between every other pulse of the target signal. The use of slower, interleaved pulses (Fig. S3B) in this concurrent, call-like sound was designed to reduce the possible influence of both energetic masking in the modulation domain (*45*) and non-simultaneous energetic masking (*46*) while retaining biological relevance: other syntopically breeding frog species produce calls with relatively slower pulse rates (*42*), including the closely-related tetraploid gray treefrog (*H. versicolor*), whose calls are generally unattractive to female *H. chrysoscelis* (*40, 47*). A separate “noise condition” (Fig. S3C) was designed to control for the possibility of energetic masking despite target signals and concurrent pulsatile sounds being well separated in spectral and modulation frequency. In this condition, the concurrent sound was a gated, bandlimited noise burst with the same duration, root-mean-square amplitude, and long-term frequency spectrum as the pulsatile concurrent sound but lacking its pulsatile envelope. Because it lacked temporal modulations, this noise condition was not expected to elicit informational masking.

#### Experiment 2: Impact of Informational Masking on Call Recognition

Male Cope’s gray treefrogs produce advertisement calls that have between 12 and 43 pulses (*41*), and females recognize calls provided they exceed, on average, about 7 or 8 pulses in length (range: 4-16 pulses) (*48*). In Experiment 2, we investigated whether informational masking impairs call recognition by testing subjects with repeating target signals that varied in pulse number (up to 20 pulses) across different trials (pulse number was fixed within a trial). Each subject was tested in the control condition and in either the pulsatile or the noise condition (Fig. 2A,C, Fig. S3). Across subjects, both target signals (1.25 kHz and 2.5 kHz) and all three noise conditions were replicated factorially at SNRs of -12 dB, -6 dB, and 0 dB (or equivalent amplitudes in the control condition).

**Fig. 2.**
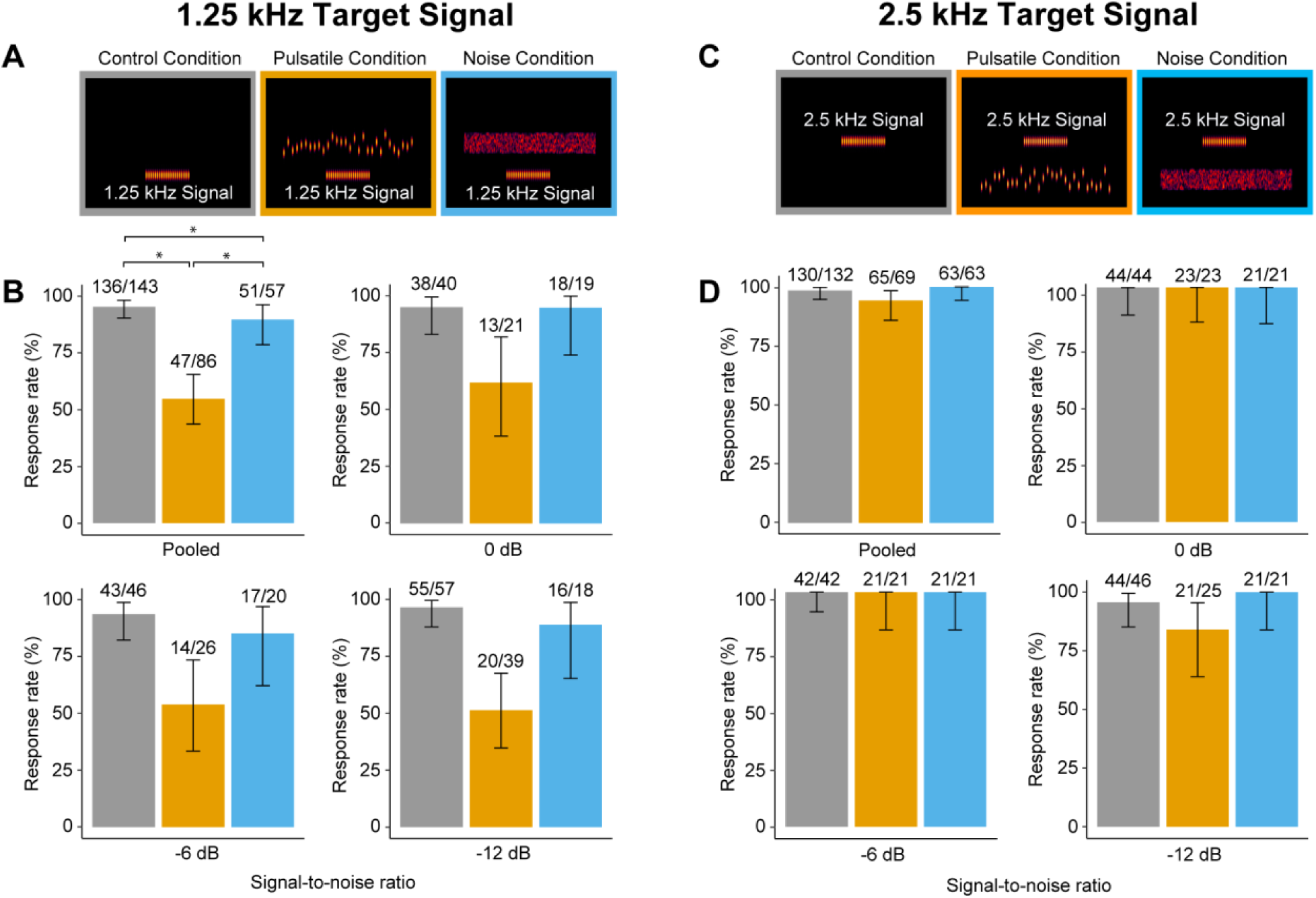
Experiment 2: Temporally modulated concurrent sounds constrained call recognition under some conditions. (**A**) Spectrograms illustrating the 1.25 kHz signal with 20 pulses in the control (*gray*), pulsatile (*orange*), and noise (*blue*) conditions. The sound pressure level of the concurrent sound was set to 75 dB SPL (LC_eq_ re 20 µPa) and that of the target signal was varied to achieve the nominal signal-to-noise ratio (SNR, or corresponding sound pressure level in the control condition. (**B**) Response rate showing the percentages (± 95% exact binomial confidence intervals) of subjects that responded to 1.25 kHz signals having 20 or fewer pulses pooled over all SNRs and separately for SNRs of -12 dB, -6 dB, and 0 dB. Fractions show the number of tests in which a subject responded over the total number of tests conducted (for pooled data) or the number of subjects that responded over the number of subjects tested (for separate SNRs). Neither SNR (χ^2^ = 1.30, P = 0.520) nor the interaction of condition and SNR (χ^2^ = 1.08, P = 0.900) had significant effects on response rates. In the pooled data, subjects were significantly (\**P* < 0.05) less likely to respond in the pulsatile condition than in the control and noise conditions, which also differed significantly. (**C**) Spectrograms illustrating the 2.5 kHz signal with 20 pulses in the control (*gray*), pulsatile (*orange*), and noise (*blue*) conditions. (**D**) Response rates as in (**B**) for 2.5 kHz signals; these data were not analyzed statistically due to the high and nearly uniform response rates across conditions.

The results revealed a pronounced effect of concurrent pulsatile sounds on the ability of females to recognize calls. Nearly all subjects (95% pooled across all SNRs) responded to the 1.25 kHz target signal in the control conditions (Fig. 2B), demonstrating robust recognition of signals with 20 or fewer pulses in the absence of concurrent sounds. A high percentage of subjects (89% pooled across all SNRs) also responded to the 1.25 kHz target signal in the noise condition (Fig. 2B), although at a slightly lower rate than in the control condition (89% vs. 95%; GEE: β = -0.97, SE = 0.456, *P* = 0.032), confirming that spectrally remote bandpass noise lacking temporal structure had only modest effects on response rates. In stark contrast, only about half of subjects (55% pooled across all SNRs) responded to the 1.25 kHz target signal in the pulsatile condition (Fig. 2B); the remainder failed to respond at all. Overall, response rates in the pulsatile condition were significantly lower than response rates in both the control (GEE: β = -2.72, SE = 0.406, *P* < 0.001) and the noise condition (GEE: β = -1.74, SE = 0.421, *P* < 0.001). Similar to results from Experiment 1, reduced response rates in the pulsatile condition were broadly consistent across a 12-dB range of SNRs (Fig. 2B).

This effect on response rates depended strongly on the frequency of the target signal (χ^2^ = 16.5, *P* < 0.001). In contrast to responses to the 1.25kHz target signal, recognition of 2.5 kHz target signals was largely unaffected by concurrent sounds. Nearly all subjects (84% to 100%) responded to a 2.5 kHz target signal having 20 or fewer pulses across all three conditions and across all three SNRs (Fig. 2D). Data for the 2.5 kHz target signal could not be analyzed statistically due to limited variation in response rates, which were high and nearly uniform at or close to 100% across conditions and SNRs.

For females that responded to a target signal in both the control and a concurrent sound condition, we also analyzed pulse-number thresholds (minimum number of pulses required to elicit phonotaxis) and response latencies. Thresholds in the pulsatile condition were occasionally elevated, but these effects were limited and inconsistent across SNRs (see Supplementary Information, Fig. S4, Table S2). Latencies did not vary across conditions in responsive females (see Supplementary Information, Fig. S4, Table S3)

Together, results from Experiment 2 are consistent with the hypothesis that informational masking interferes with the ability of females to recognize conspecific signals. A temporally structured sound at a remote frequency transduced primarily by a separate auditory end organ caused substantial impairments in call recognition, as evidenced by significantly reduced responses to the 1.25 kHz signal in the pulsatile condition. We address the asymmetry observed for different target frequencies in the Discussion. Overall, these results suggest females experienced informational masking in the pulsatile condition. Two alternative explanations can be ruled out. First, energetic masking is an unlikely explanation for reduced performance in the pulsatile condition for the following reasons. Compared with pulses in the target signal, those in the pulsatile sound occurred (i) at remote frequencies to which the sensory papillae transducing the target signal are not very sensitive, (ii) at different times (i.e., interleaved), and (iii) at half the pulse rate. Indeed, pulses in the concurrent sounds were typically an octave or more apart from target pulses in both the frequency and modulation domains. Energetic masking would generally not be expected under these conditions (*54*). In addition, the bandlimited noise, which had the same long-term frequency spectrum as the pulsatile sound but lacked temporal structure, had effects that were either negligible or small and inconsistent in how performance was affected. Second, the observed impairment in call recognition cannot be explained by any inherently aversive effects of the concurrent sounds. Neither of the concurrent sounds was aversive on its own (Fig. S2B), a result consistent with earlier studies showing that pulse trains simulating heterospecific frogs do not repel female gray treefrogs (*55*), nor do they avoid noise or inherently prefer stimuli that are quieter (*56*) or less complex (*57*).

#### Experiment 3: Impact of Informational Masking on Call Discrimination

In Experiment 3, we investigated whether informational masking interferes with the preference of females for longer calls. We conducted a series of four, two-alternative choice tests (Fig. S1B) in which subjects were allowed to choose between two alternating target signals that differed by 25% in pulse number (8 vs 10, 12 vs 15, 16 vs 20, 24 vs 30 pulses) but were otherwise identical. Most target signals were designed to have below-average pulse numbers because females discriminate more strongly at this end of the pulse-number distribution (*58*). Each choice test was again replicated factorially using both signal frequencies (1.25 kHz or 2.5 kHz) and in the control, pulsatile, and noise conditions (Fig. 3A, F). The SNR was -6 dB for all tests.

**Fig. 3.**
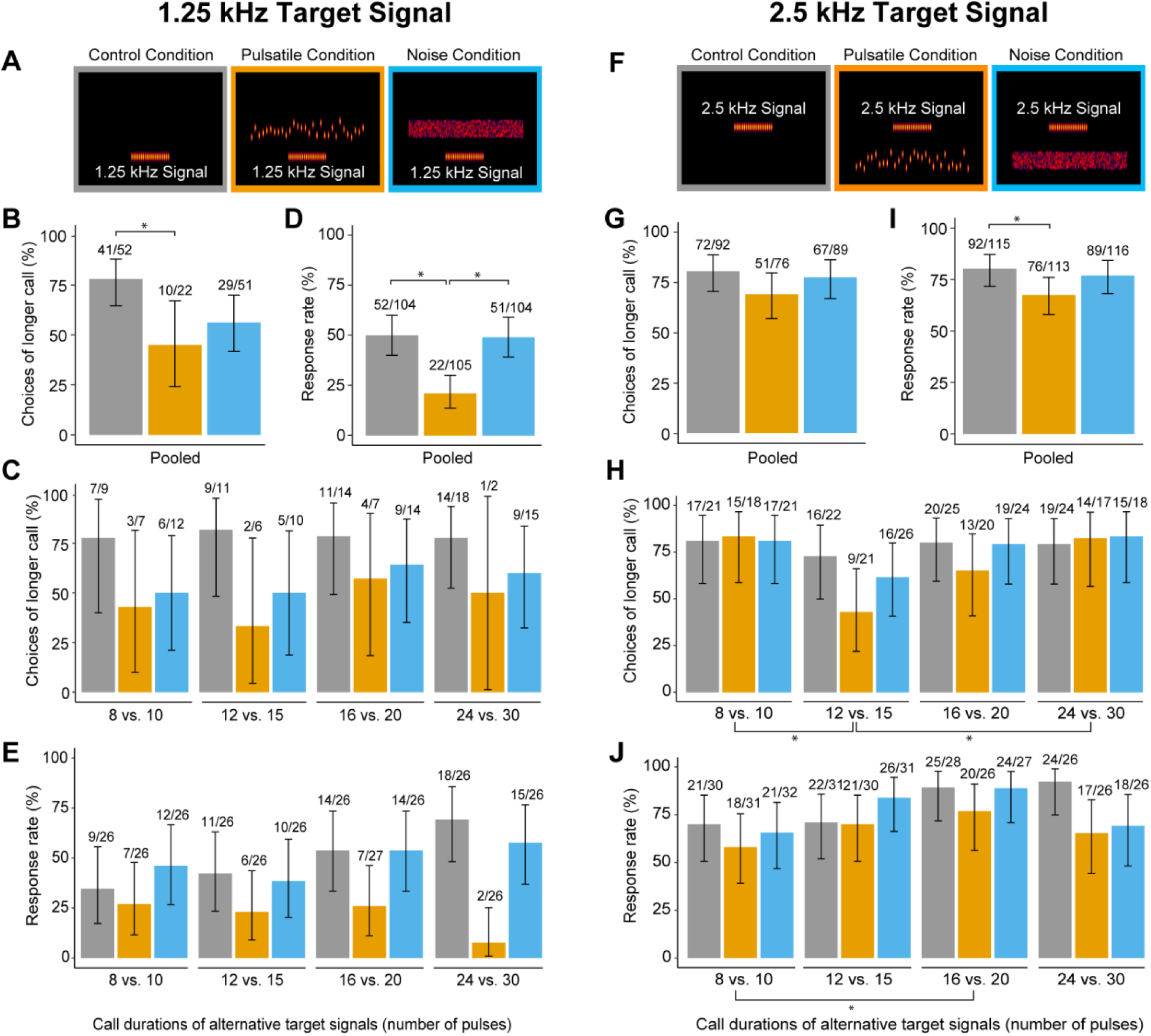
Experiment 3: Temporally modulated concurrent sounds constrained call recognition but had limited impacts on discrimination. (**A, F**) Spectrograms illustrating, respectively, the 1.25 kHz and 2.5 kHz target signals with 20 pulses in the control (*gray*), pulsatile (*orange*), and noise (*blue*) conditions. The sound pressure level of target signals was set to 69 dB (LCF re 20 µPa and that of the noise was set to 75 dB SPL (LC_eq_ re 20 µPa) to achieve a signal-to-noise ratio (SNR) of -6 dB. In separate tests, subjects were given a choice between a shorter alternative having either 8, 12, 16, or 24 pulses and a longer alternative having 25% more pulses, corresponding to pulse numbers of 10, 15, 20, or 30 pulses, respectively. (**B, G**) Preferences in two-alternative choice tests showing the percentages (± 95% exact binomial confidence intervals) of subjects that responded and chose the longer call alternative, pooled across choice tests. No significant effect of interaction between condition and pulse number was found on the proportion choosing the longer call (1.25 kHz signal: χ^2^ = 0.92, *P* = 0.989; 2.5 kHz signal: χ^2^ = 2.94, *P* = 0.816). Pulse number had no effect on preferences for the 1.25 kHz signal (χ^2^ = 0.899, *P* = 0.826) but had a significant effect for the 2.5 kHz signal (χ^2^ = 13.48, *P* = 0.004) In (**B**), subjects were significantly (\**P* < 0.05) less likely to choose the longer call in the pulsatile condition compared with the control condition (see text for details). (**C, H**) Preferences for the longer call shown separately for each choice test. Subjects were significantly (\**P* < 0.05) less likely to choose the longer call when the target signal frequency was 2.5 kHz (**H**) and the shorter alternative had 12 pulses compared with 8 pulses or 24 pulses. (**D, I**) Response rates showing the percentages (± 95% exact binomial confidence intervals) of subjects pooled across tests that responded by making a choice. There was no significant effect of interaction between condition (control, pulsatile, noise) and the number of pulses in the two alternative calls on response rates for target signals of either frequency (1.25 kHz signal: χ^2^ = 10.02, *P* = 0.124; 2.5 kHz signal: χ^2^ = 9.46, *P* = 0.150). Pulse number had no effect on response rates for the 1.25 kHz signal χ^2^ = 3.78, *P* = 0.286) but had a significant effect on responses to the 2.5 kHz signal (χ^2^ = 12.26, *P* = 0.007) In (**D**), subjects were significantly (\**P* < 0.05) less likely to respond to the 1.25 kHz target signals in the pulsatile condition compared with both the control and noise conditions (see text for details). In (**I**), subjects were significantly (\**P* < 0.05) less likely to respond to the 2.5 kHz signal in the pulsatile condition compared with the control condition after accounting for differences due to pulse number (see text for details). (**E, J**) Response rates shown separately for each choice test. Subjects were significantly (\**P* < 0.05) less likely to respond by making a choice when the target signal frequency was 2.5 kHz (**J**) and the shorter alternative had 8 pulses compared with 16 pulses. In (**B-J**), fractions indicate: (i) the number of choice tests in which subjects chose the longer call over the total number of tests in which a choice was made (for **B** and **G);** (ii) the number of subjects that chose the longer call over the number of subjects that made a choice (for **C** and **H**); (iii) the number of choice tests in which subjects responded over the total number of tests conducted (for **D** and **I**); and (iv) the number of subjects that responded over the number of subjects tested (for **E** and **J**).

Subjects chose the longer call less often in tests with the 1.25 kHz signal compared with the 2.5 kHz signal (χ^2^ = 3.96, *P* = 0.047); therefore, we statistically analyzed the data for each target signal separately. Among subjects that chose between two 1.25 kHz target signals, preferences for the longer call were strongest in the control condition, with 79% of choices favoring the longer alternative after pooling across choice tests (Fig. 3B). This preference was significantly reduced in the pulsatile condition, where only 45% of choices favored the longer call (GEE: β = -1.50, SE = 0.588, *P* = 0.011). In the noise condition, 57% of choices favored the longer call, a value that did not differ significantly from either the pulsatile condition (GEE: β = - 0.46, SE = 0.613, *P* = 0.454) or the control condition (GEE: β = -1.04, SE = 0.520, *P* = 0.046) after correcting for multiple comparisons (Fig. 3B). Among subjects that chose between two 2.5 kHz target signals, preferences for longer call were also lowest in the pulsatile condition, where 67% of choices favored the longer call compared with control and the noise conditions, where 78% and 75% of choices, respectively, favored the longer call. However, none of these differences were statistically significant when pooled across tests (Fig. 3G). For each target signal, broadly similar results were observed across pooled data and separate choice tests (1.25 kHz: cf. Figs 3B, 3C; 2.5 kHz: cf. Figs 3G, H).

The results of these choice tests provide consistent evidence that informational masking interfered with the selectivity of female preferences for longer calls, albeit only weakly. However, this interpretation must be qualified because the sample sizes in these analyses were often small (e.g., ≤ 12 in most choice tests with the 1.25 kHz signal), and hence statistical power was often limited due to the overall low response rate, particularly to short 1.25 kHz signals (Fig. 3E). This outcome reflects an important corroboration of the pronounced and differential impacts of concurrent sounds on call *recognition* observed in Experiment 2. As in Experiment 2, females tested with the 1.25 kHz signal in Experiment 3 responded at significantly lower rates (χ^2^ = 42.6, *P* < 0.001) compared with the 2.5 kHz target signal (cf. Figs 3E, J). For the 1.25 kHz target signals, response rates (Fig. 3D) were only 21% (pooled across all choice tests) in the pulsatile condition, significantly lower than both the control (50%; GEE: β = -1.36, SE = 0.385, *P* < 0.001) and the noise condition (49%; GEE: β = -1.32, SE = 0.385, *P* = 0.001), which did not differ from each other (GEE: β = 0.04, SE = 0.264, *P* = 0.884). This reduced recognition (response rate), relation to that for the pulsatile stimulus in Experiment #2, is consistent with previous findings that shorter duration pulse trains are more difficult to recognize (*48, 59*). Broadly similar, but less pronounced, impacts on response rate were also observed with the 2.5 kHz signal. Response rates for the 2.5 kHz signal (Fig. 3I) were significantly lower in the pulsatile condition (67%, pooled across all choice tests) compared with control condition (80%; GEE: β = -0.68, SE = 0.263, *P* = 0.010). Response rates in the noise condition (77%) were intermediate and more similar to the control condition (Fig. 3I), but did not differ from either the control condition (GEE: β = 0.20, SE = 0.304, *P* = 0.519) or the pulsatile condition (GEE: β = -0.48, SE = 0.343, *P* = 0.160).

Results from this experiment corroborate the findings of Experiments 1 and 2, showing that temporally modulated concurrent sounds, even at remote frequencies, strongly impair recognition of signals simulating male calls, particularly when the target signal was 1.25 kHz. Because discrimination requires successful recognition, the strong impact on recognition constrained our ability to fully evaluate pulse-number preferences for the 1.25 kHz signal. Among females that did recognize a signal, preferences for longer calls tended to be lowest in the pulsatile condition, although band-limited noise also caused reductions in preferences, suggesting that the effects of the pulsatile sound on discrimination might be more limited. Together, these results demonstrate that informational masking can substantially limit a female’s ability to make appropriate mate-choice decisions, most notably by impairing their ability to recognize and respond to calling males in natural breeding choruses.

## Discussion

Temporal amplitude modulations are crucial for encoding rhythmic and prosodic information that enables the perception and differentiation of speech and music in humans and communicative signals in nonhuman animals (*36*). This study provides direct behavioral evidence that informational masking – arising from similarity in the temporal modulation structure of signals and concurrent sounds – can disrupt vocal communication in a nonhuman animal. The finding raises important considerations for understanding the evolution and mechanisms of animal communication.

From an evolutionary perspective, the present study establishes that informational masking can constrain the recognition of vocal signals in the behavioral context of choosing a mate, which is one of the most evolutionarily consequential decisions many animals will ever make. Thus, informational masking is not a phenomenon restricted to human hearing and speech communication but should instead be considered a general acoustic communication problem shared by humans and other animals, particularly those that also communicate in noisy social settings. This conclusion has two important implications for how we study animal communication.

First, it challenges conventional views of how noise impacts nonhuman animals. The field has focused on energetic masking and the traditional filterbank model (*3, 29*) and has largely neglected the potential roles of informational masking in animal communication. While the potential for informational masking to impair communication in nonhuman animals has been recognized (*3, 29*), previous experimental studies have not investigated its impact on intraspecific communication between signalers and receivers. A few psychoacoustic studies have investigated informational masking and related phenomena using artificial sounds with nonhuman subjects (*28, 60, 61*), and a few ecological studies have explored the idea that noise causes animals to shift their attention away from behaviorally important tasks such as foraging and predator detection (*62, 63*). To our knowledge, however, all such studies have been conducted largely outside the naturalistic contexts of sexual or social communication.

Second, it serves to emphasize that informational masking of communication sounds may act as a potentially potent source of selection shaping the evolution of signals, signaling behaviors, and receiver mechanisms. Both acoustic communication and key mechanisms for hearing and receiving communication sounds have evolved independently numerous times in both invertebrates and vertebrates (*64–67*). Hence, undiscovered diversity may exist in how nonhuman animals are adapted to cope with problems of informational masking. By demonstrating the potential for informational masking to interfere with vocal signal perception, this study thus opens new avenues for future comparative research to further advance our understanding of animal hearing and acoustic communication.

From a mechanistic perspective, informational masking in humans is typically attributed to failures of auditory stream segregation or attentional constraints (*20, 26*). A failure of stream segregation is unlikely to explain our results because signals and pulsatile concurrent sounds were separated in frequency, time, or both. Although signals and concurrent sounds in our experiments originated from the same spatial location, strong spectral and temporal separation provides potent cues promoting successful stream segregation in humans and other animals (*68, 69*), including gray treefrogs (*42, 70*). Instead, our results more likely reflect constraints on bottom-up processes that contribute to the attentional salience of a signal. Bottom-up attention involves stimulus-driven mechanisms by which information is processed involuntarily based on salient stimulus features (*71, 72*). In humans, temporal modulation in competing sounds is a salient feature that interferes with bottom-up mechanisms for the temporal processing of speech (*26*) and other sounds (*45*). We found broadly similar effects in gray treefrogs: temporal modulations in concurrent sounds interfered with recognition of a temporally modulated signal. We hypothesize that these impacts resulted from disruption of bottom-up, stimulus-driven computations in the amphibian homolog of the inferior colliculus (IC).

As a key auditory processing and audio-motor integration area of the vertebrate brain, the IC functions in mediating acoustically guided behaviors with short latencies, such as responding to prey, predators, or conspecifics (*73*). In frogs, so-called “interval counting” neurons in the IC are critical for processing the temporal structure of biological sounds. In the absence of concurrent sound, interval counting neurons exhibit robust selectivity for the temporal features of conspecific calls based on bottom-up, stimulus-driven integration of time-dependent excitation and inhibition (*74, 75*). In gray treefrogs, interval counting neurons exhibit selectivity for the same temporal features of conspecific calls that determine the behavioral salience of a sound (*48, 76–78*). We suspect informational masking arises when bottom-up, stimulus-driven computations for processing temporal information in signals are disrupted by concurrent sounds. An impact on bottom-up attention may also explain the dependence of informational masking on target signal frequency in Experiments 2 and 3. Informational masking was more pronounced when concurrent pulsatile sounds fell in the range of the BP, the inner ear papilla tuned to higher frequencies. Although Cope’s gray treefrogs can process temporal information transduced by either the AP or BP, there is a frequency-dependent bias toward sounds that stimulate the higher frequency range of the BP (*51, 52*). Such bias may have contributed to the greater susceptibility to informational masking observed using concurrent pulsatile sounds that stimulated the BP and target signals that stimulated the AP (1.25 kHz).

In summary, we suggest that informational masking is an important but overlooked form of selection potentially acting on acoustic communication in a diversity of animals. Breakdowns of bottom-up attention resulting from disruptions to computations underlying temporal processing by the central nervous system may be a particularly important form of informational masking common to frogs, humans, and many other vertebrate and invertebrate animals. Additional research is needed to identify the diversity of ways informational masking may constrain animal communication, particularly in species that make evolutionarily consequential decisions in noisy social environments. It will be particularly important to determine how such constraints may, in turn, influence the evolution of signal design and the mechanisms underlying signaling behavior and signal perception. Given the growing awareness of the negative impacts of anthropogenic noise on animals (*29, 79*), it will also become necessary to consider the impacts of both energetic and informational forms of auditory masking on animal health and welfare.

## Methods

### Subjects

In total, 516 females of *H. chrysoscelis* were collected and tested in our experiments and released at their collection site after the completion of testing. All subjects were wild-caught gravid females collected by hand in amplexus at night (2200-0100 h) between mid-May and early July in 2018, 2019, 2021, 2022, 2023, and 2024 from wetlands at the Carver Park Reserve (Carver County, MN, USA), Richardson Nature Center (Hennepin County, MN, USA), and Tamarack Nature Center (Ramsey County, MN, USA). Pairs were collected in small plastic containers and returned to the laboratory on the St. Paul campus of the University of Minnesota, where phonotaxis experiments were conducted. Each pair was provided with aged tap water and maintained at approximately 4°C for up to 72 hours to delay oviposition. This is a common procedure used with temperate zone frogs that delays oviposition and extends a female’s responsiveness in behavioral tests (*43*). All animal procedures were approved by the University of Minnesota’s Institutional Animal Care and Use Committee (1701-34456A, 2001-37746A, 2301-40692A).

### General protocol

Approximately 30 minutes before testing, each collected pair was transferred to a temperature-controlled incubator so that their body temperature could reach 20 ± 1°C. All experiments were conducted at this temperature because female preferences are often temperature-dependent (*43*), and 20°C is close to the average nighttime temperature at the local field sites during the gray treefrog breeding season (*41*). For testing, females were separated from their chosen mate, tested, then reunited with their mate in the incubator for a brief (typically 5-10 min) timeout between consecutive tests.

We performed two types of phonotaxis tests – no choice tests (Fig. S1A) and two-alternative choice tests (Fig. S1B) – that are well established in studies of acoustically guided behavior in frogs and other animals (*38, 43*). Both types involved repeatedly testing subjects across a series of phonotaxis tests. In no-choice tests (Experiments 1 and 2), the stimulus was presented through a single speaker, and we determined whether the subject exhibited phonotaxis toward the speaker. In choice tests (Experiment 3), we gave females a choice between two stimuli broadcast from two speakers in an alternating fashion, with equal periods of silence occurring before and after each stimulus. The stimulus assigned to each speaker in a choice test was determined randomly. There is little evidence that repeat phonotaxis testing in female frogs has carryover effects (*80*).

Target signals and concurrent sounds were synthesized in MATLAB R2018a (Mathworks, Natick, MA, USA) at a sampling rate of 44.1 kHz (16 bit). In all phonotaxis tests, target signals were repeated at a rate of 11 calls/min, which approximates the average call rate in local populations at 20°C (*41*). The pulses of all target signals and concurrent pulsatile sounds had the identical gross temporal structure and were modeled after the average pulse shape in local populations of *H. chrysoscelis* and (10 ms duration, 3.1-ms inverse exponential rise time, 5.4-ms exponential fall time).

We conducted phonotaxis tests using a circular arena (Fig. S1C; 2-m diameter, 60-cm height) inside a custom-built, temperature-controlled (20 ± 1°C), semi-anechoic chamber (length x width x height: 2.8 x 2.3 x 2.1 m; Industrial Acoustics Company, IAC, North Aurora, IL, USA). The chamber walls and ceiling were lined with dark gray acoustic absorber panels (IAC’s Planarchoic^TM^ system), and the floor was covered with dark gray, low-pile carpet. The arena was made from hardware cloth and covered with black fabric. On the floor outside the arena wall, we positioned two Mod1 Orb speakers (Orb Audio, Sherman Oaks, CA, USA) separated by 90°, with both speakers directed toward the center of the arena (Fig. S1C). Stimuli were broadcast using Adobe Audition (Adobe Systems Inc., San Jose, CA, USA) running on a Dell Optiplex 980 or 5050 PC (Dell Computer Corporation, Round Rock, TX, USA). We used a MOTU model 16A sound card (MOTU, Inc., Cambridge, MA, USA) interfaced with a Crown XLS 1000 High-Density Power Amplifier (Crown International, Los Angeles, CA, USA) to broadcast sounds. Sound levels were determined by placing a Bruël and Kjær Type 4950 microphone connected to a Bruël and Kjær Type 2250-L sound level meter (Bruël and Kjær, Nærum, Denmark) 1 m away from the speaker at the approximate position of a subject’s head at the start of a phonotaxis test. At this position, the frequency response of the playback system was flat (± 2 dB) between 0.7 kHz and 2.8 kHz.

At the beginning of a phonotaxis test, an individual subject was placed in a small acoustically transparent release cage (9-cm diameter, 2-cm height) located on the center of the arena floor. We allowed the subject to acclimate in the cage for 30 sec, after which we started playback of the target signal either in the absence of concurrent sound (i.e., the control conditions) or with the designated concurrent sound. At this time, we remotely lifted the lid of the release cage using a rope-and-pulley system and allowed the subject to move freely in the arena. Subject behavior was observed under infrared light and scored in real-time by an observer outside the chamber by means of a video camera and monitor. A ”response” was scored if the subject entered a response zone consisting of a 10-cm semi-circle in front of a playback speaker. We scored a “no response” if the subject did not exit the release cage within 3 min, did not enter a response zone within 5 min, or first made contact with the arena wall in the semi-circle opposite the speaker in a no-choice test or in the quadrant opposite that formed by the two speakers in a two-alternative choice test.

### Experimental designs

Experiment 1 (*n* = 130) consisted of no-choice tests (Fig. S1A) in which a target signal with two spectral components at 1.25 kHz and 2.5 kHz (30 pulses, 50 pulses/s, 20-ms pulse period, 50% pulse duty cycle) was presented in the absence of concurrent sound (control condition) or in the presence of a continuous concurrent sound consisting of a 4-component harmonic complex that was either pulsatile (50 pulses/s, 20-ms pulse period, 50% pulse duty cycle) or unmodulated. The fundamental frequency (f_0_) of the harmonic complex in each concurrent sound varied between 250 Hz and 500 Hz in 2-semitone steps. To generalize our results, f_0_ was chosen randomly for each trial in the unmodulated condition and for each pulse on each trial in the pulsatile condition. Subjects were assigned randomly to the control (*n* = 25), pulsatile (*n* = 25), and unmodulated conditions (*n* = 25), and each subject was tested at five SNRs (-18, -12, -6, 0, and +6 dB; or equivalent signal levels in the control conditions) using a within-subjects design, with SNR order determined randomly. The subject’s response was coded as binary outcome (1= response, 0 = no response). If at any time during testing, a subject failed to respond in three consecutive tests, they were next tested with a salient conspecific call to ensure they remained motivated to respond. Only animals that exhibited phonotaxis in this “motivation test” were used as subjects. We conducted additional “sham” tests in which the pulsatile (*n* = 30) and unmodulated (*n* = 25) concurrent sounds were presented alone (i.e., no target signal). For these two additional tests, the response variable was the angle at which subjects first touched the wall of the arena to assess whether the concurrent sounds, by themselves, elicited positive or negative phonotaxis (Fig. S2). We conducted a motivation test before and after each sham test.

In contrast to Experiment 1, Experiments 2 and 3 presented target signals consisting of a single spectral component (1.25 kHz or 2.5 kHz) in the presence of concurrent sounds that were presented in the opposite frequency range and gated on and off with each target signal presentation. Target signals varied in the number of pulses (see below). The same concurrent sounds were used in both Experiments 2 and 3 and consisted of either a series of 30 pulses (25 pulses/s, 40-ms pulse period, 25% pulse duty cycle) or a bandlimited noise. In the pulsatile condition, the carrier frequency of each pulse in each successive concurrent sound was randomly chosen from among 13 possible frequencies spanning a range of 0.7 kHz. For the 2.5 kHz target signal (BP range), these frequencies were spaced one semitone apart between 0.7 kHz and 1.4 kHz (0.7, 0.742, 0.786, 0.832, 0.882, 0.934, 0.99, 1.049, 1.111, 1.177, 1.247, 1.321, and 1.4 kHz) and were chosen to fall within the frequency range of the AP but below the minimum frequency to which the BP responds (*44*). For the 1.25 kHz signal (AP range), the 13 possible frequencies had the same absolute spacing (in kHz) but were shifted upward in frequency by 1.4 kHz so that they fell between 2.1 and 2.8 kHz (2.1, 2.142, 2.186, 2.232, 2.282, 2.334, 2.390, 2.449, 2.511, 2.577, 2.647, 2.721, and 2.8 kHz) and were above the maximum frequency to which the AP responds (*44*). The bandlimited noise used in the noise conditions of Experiments 2 and 3 had the same duration and bandwidth (0.7 kHz to 1.4 kHz or 2.1 kHz to 2.8 kHz) as the corresponding pulsatile sound but lacked a pulsatile temporal envelope. In both conditions, each presentation of a target signal was temporally centered in a concurrent sound. Each separate rendition of a concurrent pulsatile sound or noise burst was randomized: for pulsatile sounds, the carrier frequency of individual pulses was randomly selected, and for the bandlimited noise, each burst consisted of the newly generated noise waveform with the same frequency spectrum as the corresponding pulsatile sound. As a result, the same combination of target signal and concurrent sound never repeated in the same test of a given subject.

In Experiment 2 (*n* = 275), we assessed signal recognition using no-choice tests (Fig. S1A) and an adaptive tracking procedure to measure pulse number thresholds following (*48*). Briefly, for each subject, we determined a pulse number threshold in the control condition and in one of the two concurrent sound conditions, with order determined randomly. Subjects were assigned randomly to a concurrent sound condition (pulsatile or noise), a target signal frequency (1.25 kHz or 2.5 kHz), and one of three SNRs (-12 dB, -6 dB, or 0 dB). Within each condition, the target signal had 8 pulses on the subject’s first test. Depending on the subject’s response, we either decreased or increased the number of pulses in the target signal in subsequent tests until the subject’s behavior changed (see additional details in Gupta et al. 2021). If subjects did not respond to a target signal with 20 or fewer pulses, its pulse number threshold was not determined. Restricting the maximum number of pulses to 20 allowed us to ensure we could measure two pulse number thresholds per subject, which typically do not respond in more than about 15 to 20 phonotaxis tests before losing response motivation. After determining a subject’s first pulse number threshold we immediately restarted the adaptive tracking procedure to determine their second threshold. Animals that failed to respond in three consecutive tests were given a motivation test; only animals that exhibited phonotaxis were used as subjects. For each subject in each condition, pulse number threshold was calculated by averaging the lowest pulse number that elicited a response and the highest pulse number that did not. We determined response rate in each condition by coding each subject’s response as binary (1 = responded to 20 or fewer pulses; 0 = no response). We determined response latency in each condition as the time required for each subject to respond to the target signal having the lowest number of pulses that elicited a response. Pulse number thresholds were determined similarly in the control and concurrent sound conditions with the following exception. Prior to commencing the adaptive tracking procedure in the concurrent sound conditions, subjects experienced a sham test in which the concurrent sound was presented alone without the target signal. To obtain accurate estimates of pulse number thresholds we excluded animals that responded in the sham condition (Fig. S2).

In Experiment 3 (*n* = 111), we assessed signal discrimination using two-alternative choice tests (Fig. S1B) to measure female preferences for longer calls with more pulses. Separate groups of subjects were randomly assigned to tests with the 1.25 kHz or 2.5 kHz signal. Subjects were given a choice between a shorter target signal (with 8, 12, 16 pulses, or 24 pulses) and a longer target signal having 25% more pulses (10, 15, 20, and 30 pulses, respectively). The tests of 8 versus 10, 12 versus 15, and 16 versus 20 pulses were performed within subjects, and all three of these tests were also tested within subjects across the three conditions (control, pulsatile, noise); a separate group of subjects was tested with 24 versus 30 pulses in all three conditions. In the pulsatile and noise conditions, the two temporally adjacent target signals broadcast from separate speakers were played with the same rendition of the assigned concurrent sound; renditions varied randomly across successive pairs of the two alternating signals over the duration of the choice test. The order of choice tests was randomized for each subject. The shorter signal was always the signal that started the alternating sequence of two target signals. We recorded whether a subject responded by making a choice (1= response, 0 = no response) and if so, we scored its choice (1 = longer, 0 = shorter). We performed a motivation test after three consecutive no responses to confirm females were still motivated to respond. Subjects that lost motivation during testing (7 of 111) were not tested further, but we included their data from all choice tests completed before they stopped responding.

The sound pressure levels (SPL; LC_eq_ re 20 µPa at 1 m) of concurrent sounds were calibrated to 79 dB SPL in Experiment 1 and 75 dB SPL in Experiments 2 and 3. These levels approximate the signal and noise levels females likely experience while listening to calling males in dense choruses (*81*). Experimental SNRs for presenting target signals in the presence of concurrent sounds were achieved by varying the SPL of the target signal (LCF) across five levels in Experiment 1 (61, 67, 73, 79, and 85 dB SPL), three levels in Experiment 2 (63, 69, and 75 dB SPL), and one level in Experiment 3 (69 dB SPL). The highest signal level used (85 dB SPL) approximates a male calling at a distance of about 1 m (*82*).

### Statistical analyses

All statistical analyses were conducted using R version 4.3.1. A significance criterion of ɑ = 0.05 was used, and the Holm-Šydák correction for multiple comparisons was applied within each experiment. We analyzed phonotaxis responses using a combination of Generalized Estimating Equations (GEE) and nonparametric statistics. GEE models are an extension of generalized linear models that account for correlated or clustered data, making them well suited for behavioral experiments in which binary responses are obtained from same subjects under different stimulus conditions (*83*). All GEE models used binomial logit link functions and exchangeable correlation structures, and subject ID was always included as the clustering factor to account for repeated measures. Although Generalized Linear Mixed Models (GLMMs) can also accommodate repeated measures, they estimate random effects that capture individual-level variability. Because our primary goal was to infer population-level relationships between phonotaxis behavior and experimental factors, GEE provided more robust and interpretable estimates without the need to specify random-effect structures.

For Experiment 1 (Fig. 1), GEE models included condition (control, pulsatile, unmodulated), SNR (continuous variable), and their interaction as predictor variables. Model comparison via ANOVA indicated a significant condition × SNR interaction (χ^2^ = 161, *P* < 0.001). Therefore, the full interaction model was retained. For each condition, we tested whether response rates increased with SNR by examining the slope of response rate versus SNR in the GEE model. A significant positive slope was interpreted as performance improvement with increasing signal level, whereas a non-significant or flat slope indicated no change in response rates across SNRs. To evaluate how differences between acoustic conditions varied across SNR, we additionally fitted a GEE model with SNR treated as a categorical variable.

In Experiment 2 (Fig. 2), an initial inspection of the data revealed large effects of target signal frequency, which we confirmed by comparing two GEE models with and without signal frequency as a predictor variable for each of three response variables (response rate, pulse number threshold, and response latency). Because the effects of target signal frequency were of secondary interest, we analyzed the effects of condition and SNR separately for each signal frequency. For analyses of response rate with the 1.25 kHz signal, the initial GEE model included condition (control, pulsatile, noise), SNR (-12 dB, -6 dB, and 0 dB), and the condition × SNR interaction as predictor variables. We fitted a second GEE model in which we excluded the condition × SNR interaction. We compared the two models using ANOVA and found that they were not statistically different (χ^2^ = 1.08, *P* = 0.897). We fitted a third GEE model in which we additionally excluded SNR as a predictor variable and compared this model with the second model using ANOVA and found that they also were not statistically different (χ^2^ = 1.30, *P* = 0.521). For analyses of response rate with the 2.5 kHz signal, we could not fit GEE models because the response rate of subjects was high across all conditions, reaching 100% in seven of the nine factorial combinations of condition and SNR. The lack of variance in the data led to the problem of complete separation, producing spurious results for the main effects model.

Also for Experiment 2, we compared pulse number thresholds and response latencies for each subject that responded both in the control condition and in their assigned concurrent sound condition. This paired data allowed us to evaluate how the presence of each concurrent sound influenced responses relative to a signal alone condition in the same group of subjects. Because these data were not normally distributed and could not be successfully transformed to follow a normal distribution, we used non-parametric tests for hypothesis testing. To make pairwise comparisons, we used a paired-sample Wilcoxon tests at each SNR to compare separately the pulse number thresholds and response latencies in the control condition and in the paired condition with concurrent sound. An independent sample Wilcoxon test was used to compare pulse number thresholds and response latencies between subjects that responded in the pulsatile and noise conditions at each SNR.

For Experiment 3, we used GEE to analyze response rates and choices. For each response variable, we initially fitted models with and without target signal frequency as a predictor variable. Because signal frequency again had a significant effect and was of secondary interest, we used separate GEE models to analyze results for the 1.25 kHz and 2.5 kHz signals. We next used a parallel series of model fitting for each response variable. The first model included as predictor variables the condition (control, pulsatile, noise), the number of pulses in the shorter signal (8, 12, 16, or 24 pulses), and the condition × pulse number interaction. We compared this model with a second model in which we excluded the interaction term. We did not find any statistical difference between the two models for either signal frequency (response rates: 1.25 kHz: χ^2^ = 10.02, *P* = 0.124, 2.5 kHz: χ^2^ = 9.46, *P* = 0.150; choices: 1.25 kHz signal: χ^2^ = 0.92, *P* = 0.989; 2.5 kHz signal: χ^2^ = 2.94, *P* = 0.816). We next fitted a third model in which we additionally excluded pulse number as a predictor variable and compared the second and the third model. If pulse number had a significant effect on the outcome, we adopted the second model. Otherwise, we adopted the third model with condition as the only predictor variable.

## Acknowledgments

We thank Paloma Gonzales-Bellido, Katie Krueger, Andrew Oxenham, Annika Rupert, and Emilie Snell-Rood for feedback on the manuscript; Andrew Oxenham for insightful conversations about informational masking; Mridul Yadav for making the 3D schematic of the phonotaxis arena; all members of the Bee Lab 2018-2024 for their help collecting and testing frogs; the Minnesota Department of Natural Resources for permission to collect frogs; and the Three Rivers Park District and the Ramsey County Department of Parks and Recreation for access to field sites.

## Funding

This study was funded by the U.S. National Science Foundation (IOS-2022253 to MAB, GJR).

## Author contributions

S.G., G.J.R., and M.A.B. designed research; S.G., L.K., and L.D. performed research; S.G. analyzed data; S.G and M.A.B. wrote the manuscript; all authors edited the manuscript.

## Competing interests

The authors declare that they have no competing interests.

## Data availability

All data and code associated with this manuscript are available on Mendeley data (https://data.mendeley.com/preview/73rt9t5mcd?a=daec2bb8-1b79-4327-aa1c-bec453ecacb0).

## Supplementary Materials for

**Fig. S1.**
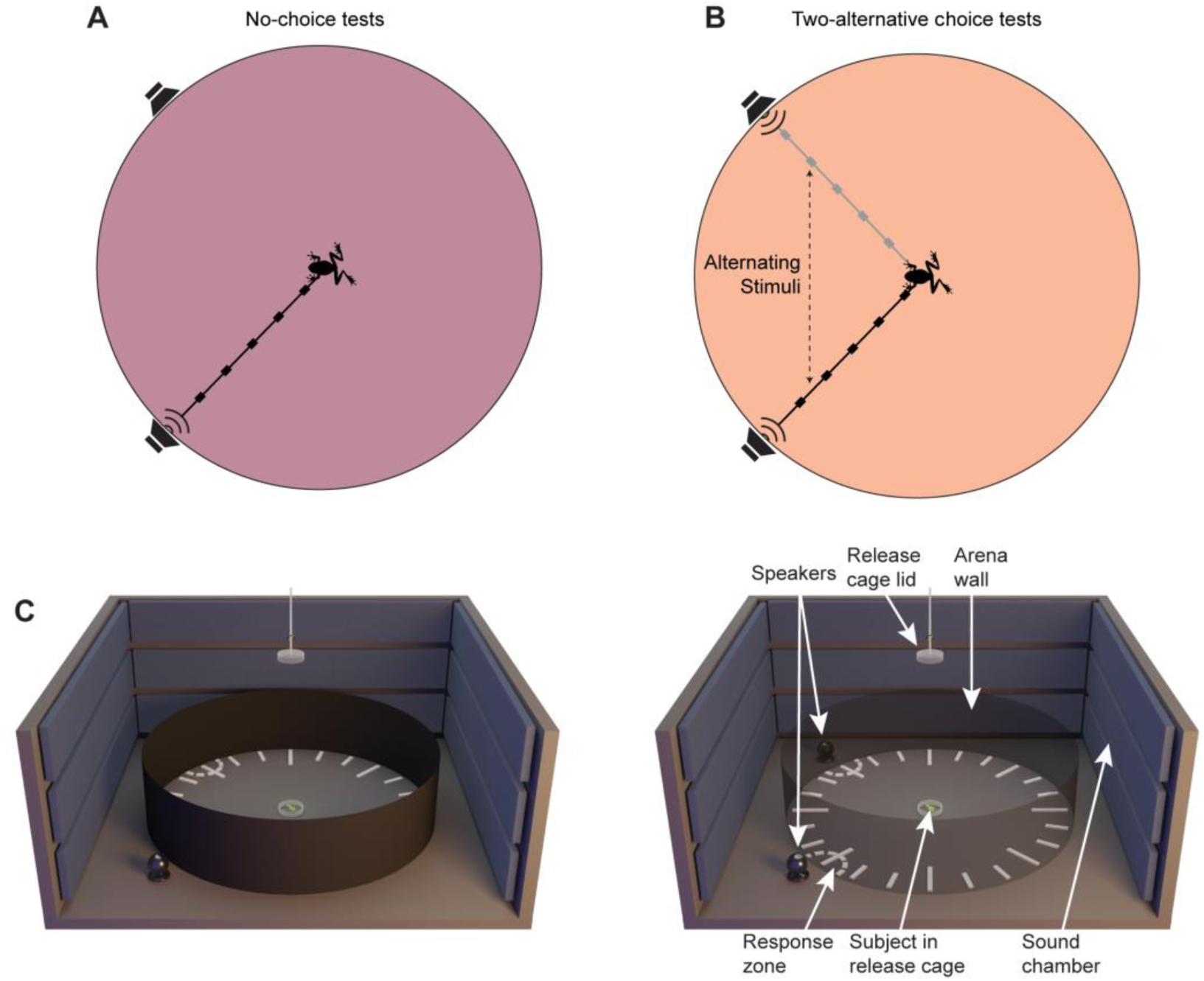
Schematic illustrations of testing protocols and experimental setup. (A,B) Overhead schematics (not to scale) of the testing arena illustrating a single stimulus, no-choice test **(A)** and a two-alternative choice test **(B)**. **(A)** In no-choice tests, a single repeated stimulus was presented from one of the two speakers separated by 90° around the test arena. **(B)** In two alternative choice tests, two sequences of repeating stimuli (e.g., a signal alone or a signal plus concurrent sound) were presented from the two speakers; stimuli in the two sequences alternated in time such that there were equal intervals of silence preceding and following each stimulus presentation. To minimize any side bias, we randomized which speaker broadcast which stimulus across subjects. Alternating stimulus sequences are illustrated by gray and black schematic waveforms. **(C)** A 3D schematic of the circular test arena used to conduct phonotaxis tests with female treefrogs shown with visually opaque walls (Left) with walls rendered transparent (Right) to facilitate visualization of both playback speakers. The test arena (2-m diameter, 60-cm height) was located inside a semi anechoic chamber and was made from hardware cloth covered in black fabric. It was acoustically transparent but visually opaque. Acoustic stimuli were broadcast from one or two speakers positioned on the floor outside the arena wall and pointed toward a subject release cage at the center of the arena. The lid of the release cage could be remotely lifted from outside the chamber to allow the subject to respond by approaching a response zone. A response was scored when the subject entered a response zone. Responses were recorded using an infrared sensitive camera mounted above the center of the test arena (not shown) and viewed in real time by one or two experimenters on a monitor located outside the arena.

**Fig. S2.**
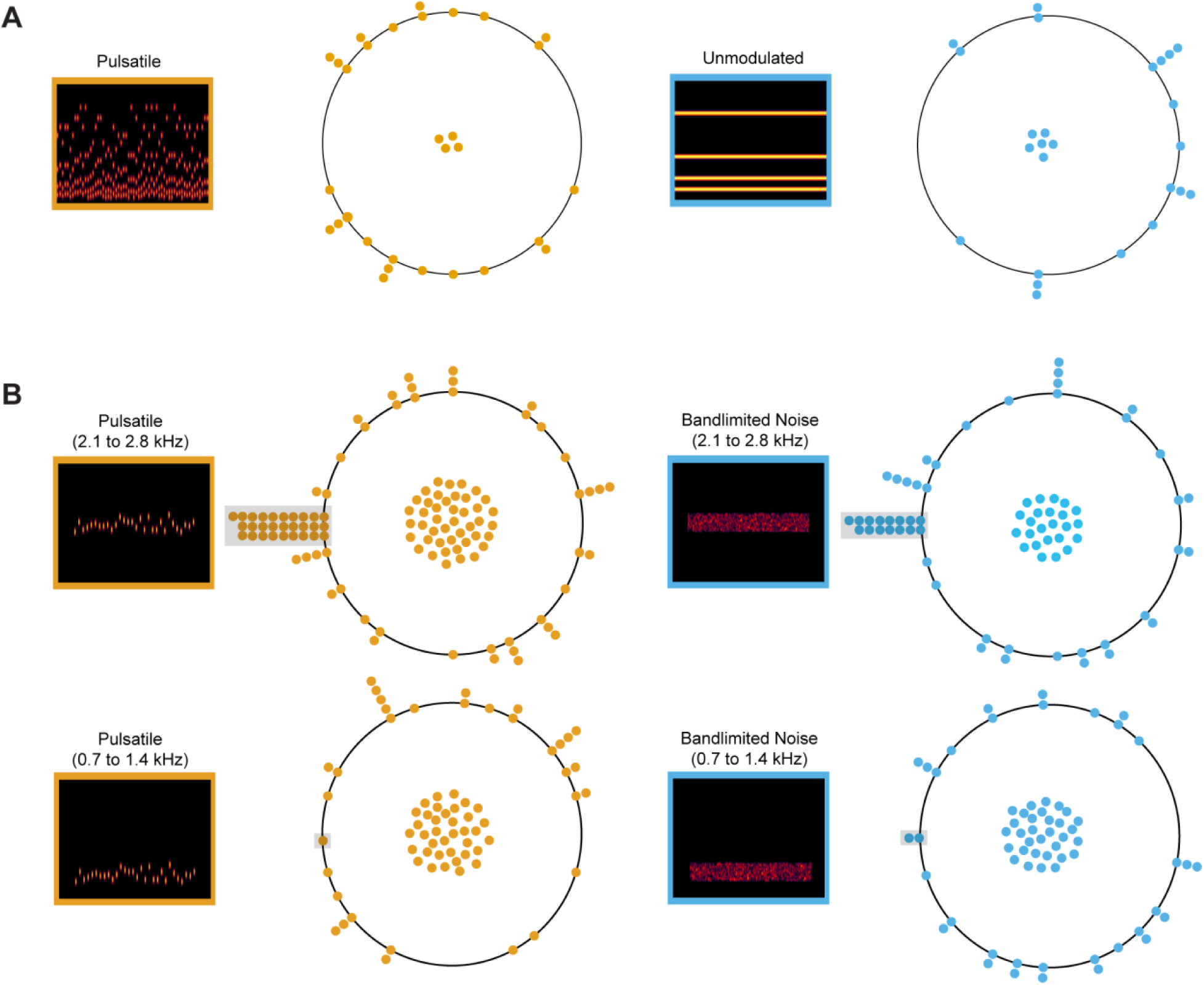
Concurrent sounds presented by themselves were not behaviorally aversive. Points on each circle show where subjects tested in sham trials (i.e., no target signal) first touched the wall of the circular test arena relative to the playback speaker positioned at 9 o’clock around the circle. Individuals that never exited the release cage are depicted as points at the center of each circle and are excluded from the circular statistics (Rayleigh tests of uniformity) reported here. Insets show spectrograms illustrating the stimulus used in each sham trial. **(A)** The pulsatile and unmodulated sounds used in Experiment 1 were not behaviorally aversive, eliciting neither positive nor negative phonotaxis (pulsatile sound: r = 0.260, *P* = 0.172; unmodulated sound: r = 0.382, *P* = 0.061). **(B)** The pulsatile sound and bandlimited noise used in Experiments 2 and 3 were also not behaviorally aversive. Some individuals exhibited positive phonotaxis (gray shaded regions) to the pulsatile sound (*n* = 28 of 114) and bandlimited noise (*n* = 15 of 72) in a frequency range (2.1 to 2.8 kHz) transduced by the basilar papilla (BP; **B**, *top row*). After excluding these individuals, the remaining subjects exhibited neither positive nor negative phonotaxis (pulsatile sound: r = 0.021, *P* = 0.981; bandlimited noise: r = 0.068, *P* = 0.861). In contrast, few subjects exhibited positive phonotaxis to the pulsatile sound (*n* = 1 of 70) and bandlimited noise (*n* = 2 of 65) in a frequency range (0.7 to 1.4 kHz) transduced by the amphibian papilla (AP; **B,** *bottom row*). After excluding these individuals, the remaining subjects exhibited neither positive nor negative phonotaxis (pulsatile sound: r = 0.286, *P* = 0.071; bandlimited noise: r = 0.131, *P* = 0.590). All individuals that exhibited positive phonotaxis were excluded Experiment 2 to avoid introducing ambiguity into estimates of pulse number thresholds. A significantly greater proportion of subjects exhibited positive phonotaxis toward stimuli presented in the BP range than in the AP range (two proportion z-test: pulsatile, χ^2^ = 15.8, *P* < 0.001; band-limited noise, χ^2^ = 8.5, *P* = 0.002). This differential response to concurrent sounds that were similar in all aspects other than their frequency range corroborates previous findings showing that high-frequency sounds are perceptually more salient than low-frequency sounds in *H. chrysoscelis*.

**Fig. S3.**
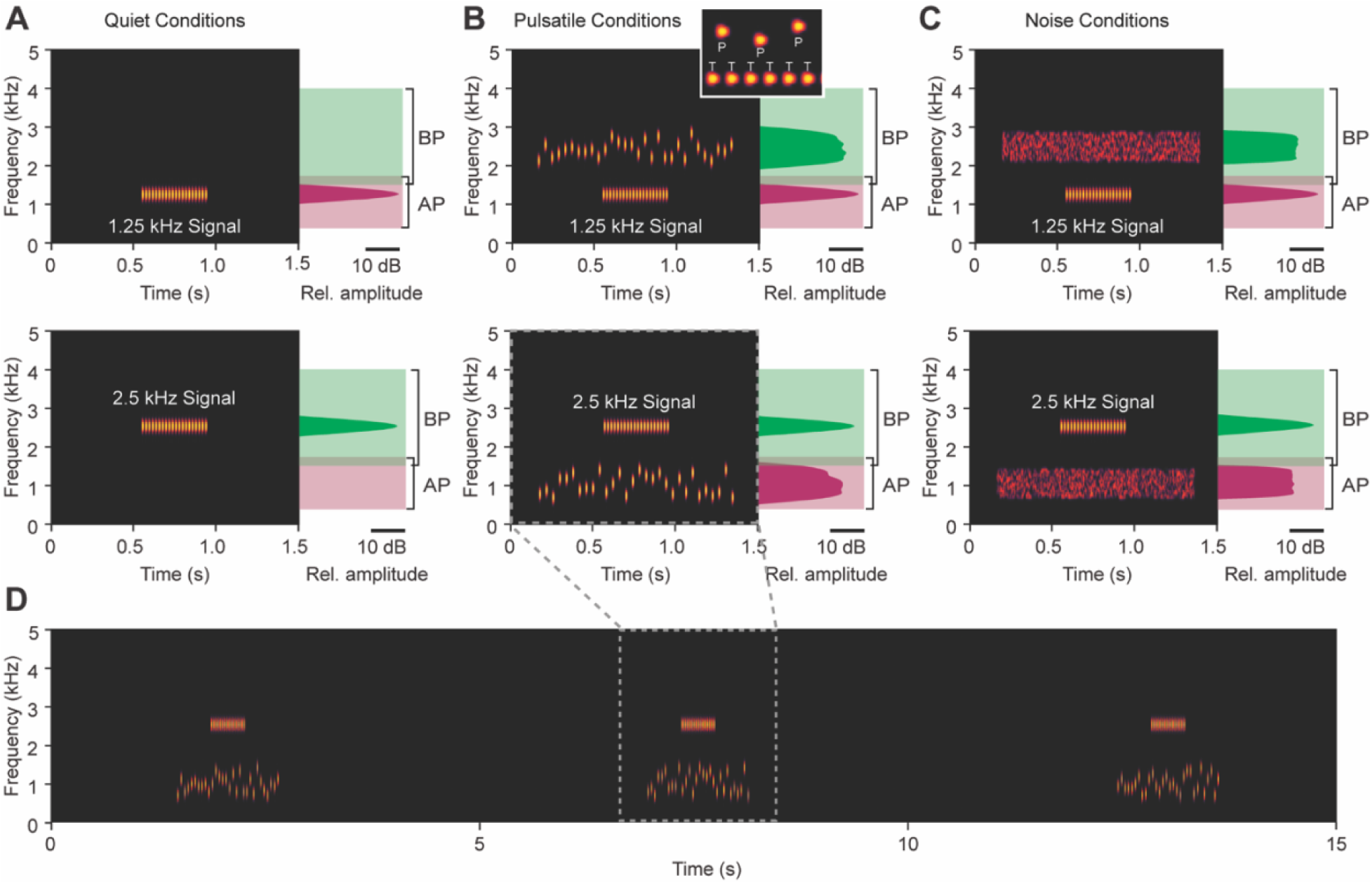
Examples of stimuli used to investigate call recognition in Experiments 2 and call discrimination in Experiment 3. The 1.25 kHz and 2.5 kHz target signals (20 pulses) are shown separately in the (**A**) control, (**B**) pulsatile, and (**C**) noise conditions. Each plot in **A-C** shows a spectrogram (frequency × time) and power spectrum (frequency × relative amplitude) of one repetition of a target signal either in control (**A**) or with its associated concurrent sound (**B, C**). The inset in (**B**) illustrates how the pulses (P) of the concurrent sound in the pulsatile condition were temporally interleaved with every other pulse in the target signal (T). Power spectra are depicted relative to the approximate frequency ranges of the basilar papilla (BP; green) and amphibian papilla (AP; pink), the two sensory papillae in the frog inner ear that transduce airborne sound frequencies. Based on recordings of the auditory brainstem response in gray treefrogs, the lower spectral component (∼1.25 kHz) is transduced primarily by the AP (range of sensitivity ∼0.4 to 1.75 kHz), and the higher component (∼2.5 kHz) is transduced primarily by the BP (range of sensitivity ∼1.5 to 4.0 kHz). As illustrated, there is minimal overlap in the tuning of the two sensory papillae. By specifying the carrier frequency of the target signal at either 1.25 kHz or 2.5 kHz and limiting concurrent sounds to the opposite frequency range, the stimuli were designed such that target signals and concurrent sounds would primarily stimulate different inner ear papillae. The concurrent sound in the pulsatile condition was a series of random-frequency pulses (25 pulses/s) that were interleaved between every other pulse in a target signal (50 pulses/s). The narrowband noise in the noise condition had the same spectral content and long-term sound pressure level as the pulsatile concurrent sound. In both conditions with concurrent sounds, target signals were temporally centered within the concurrent sound, which was gated on and off around the signal. (**D**) A 15-s segment of a stimulus using in Experiments 1 and 2. During phonotaxis tests, stimuli were repeated at a rate of 11 calls/min for up to 5 minutes or until a subject responded.

**Fig. S4.**
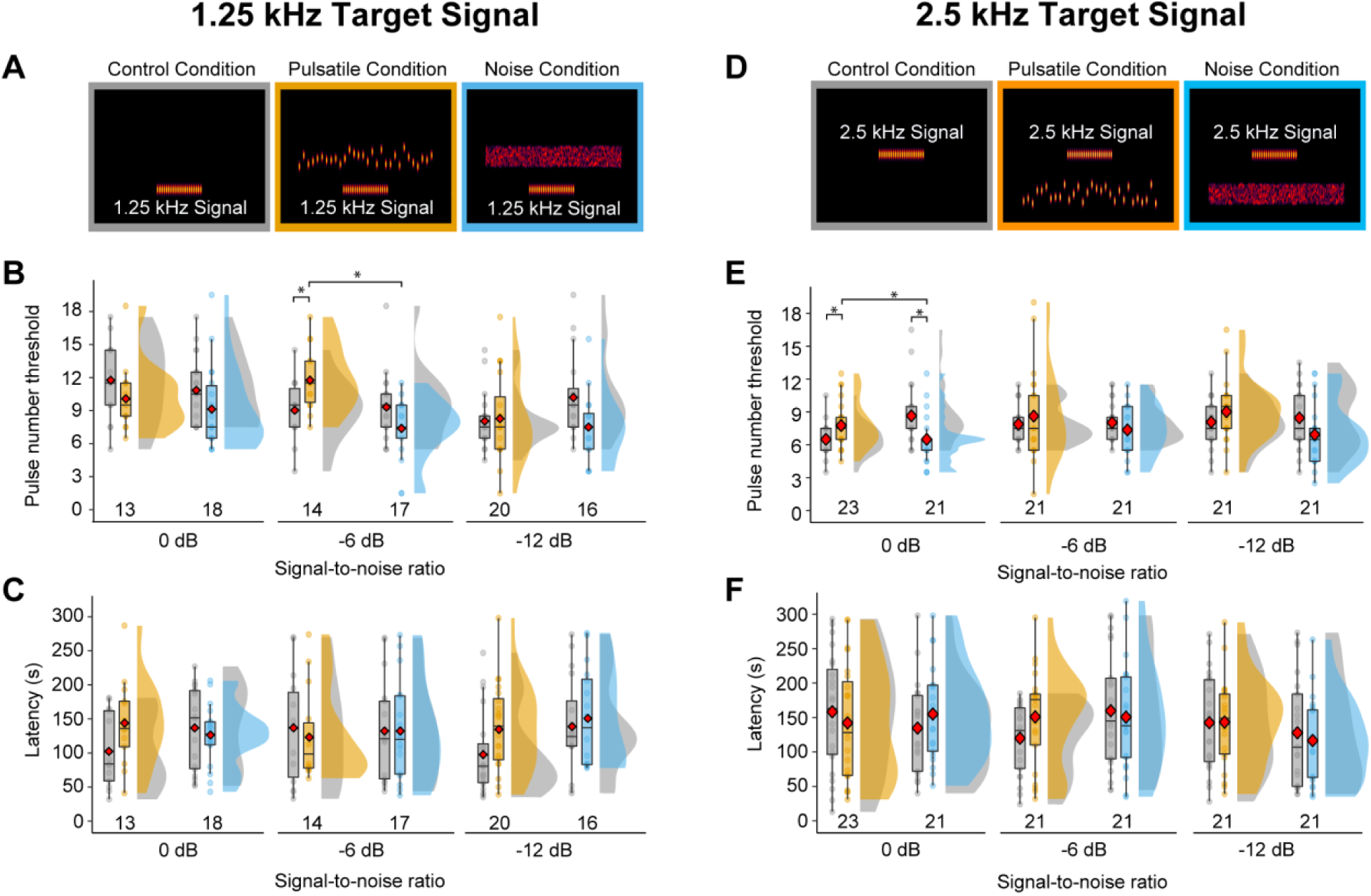
Experiment 2: Temporally modulated concurrent sounds had limited impact on pulse number thresholds and response latencies. (**A**) Spectrograms illustrating the 1.25 kHz signal with 20 pulses in the control (*gray*), pulsatile (*orange*), and noise (*blue*) conditions. (**B**) Pulse number thresholds for responding to 1.25 kHz signals measured using an adaptive tracking procedure and defined as the minimum number of pulses required to elicit positive phonotaxis. For each subject tested at each SNR, separate thresholds were determined within subjects in the control condition and in either the pulsatile condition or the noise condition. Box plots depict the first, second (median), and third quartiles, and the mean (red diamond) values; whiskers depict 1.5 times the interquartile range and points depict individual data points. The two overlapping distributions are kernel density representations of the data that are depicted in the corresponding box plots. For each SNR, the two box plots and distributions on the left show data for the control and pulsatile conditions that were paired within subjects, whereas the two box plots and distributions on the right show data for the control and noise conditions that were paired within a different group of subjects. Sample sizes are indicated at the base of each dataset. At a SNR of -6 dB, pulse number thresholds were significantly higher (\**P* < 0.05) in the pulsatile condition compared with the paired control condition and the noise condition (Table S1). (**C**) Response latencies for the 1.25 kHz signal, computed as the time to respond to the target signal having the lowest number of pulses that elicited positive phonotaxis. There were no significant differences in response latencies. Box plots and distributions as in (**B**). (**D**) Spectrograms illustrating the 2.5 kHz signal with 20 pulses in the control (*gray*), pulsatile (*orange*), and noise (*blue*) conditions. (**E**) Pulse number thresholds for responding to 2.5 kHz signals. At a SNR of 0 dB, pulse number thresholds were significantly higher in the pulsatile condition compared with both the paired control condition and the noise condition; they were significantly lower in the noise condition compared with the paired control condition (Table S2). (**F**) Response latencies for 2.5 kHz signal. There were no significant differences in response latencies (Table S3).

**Table S1.**
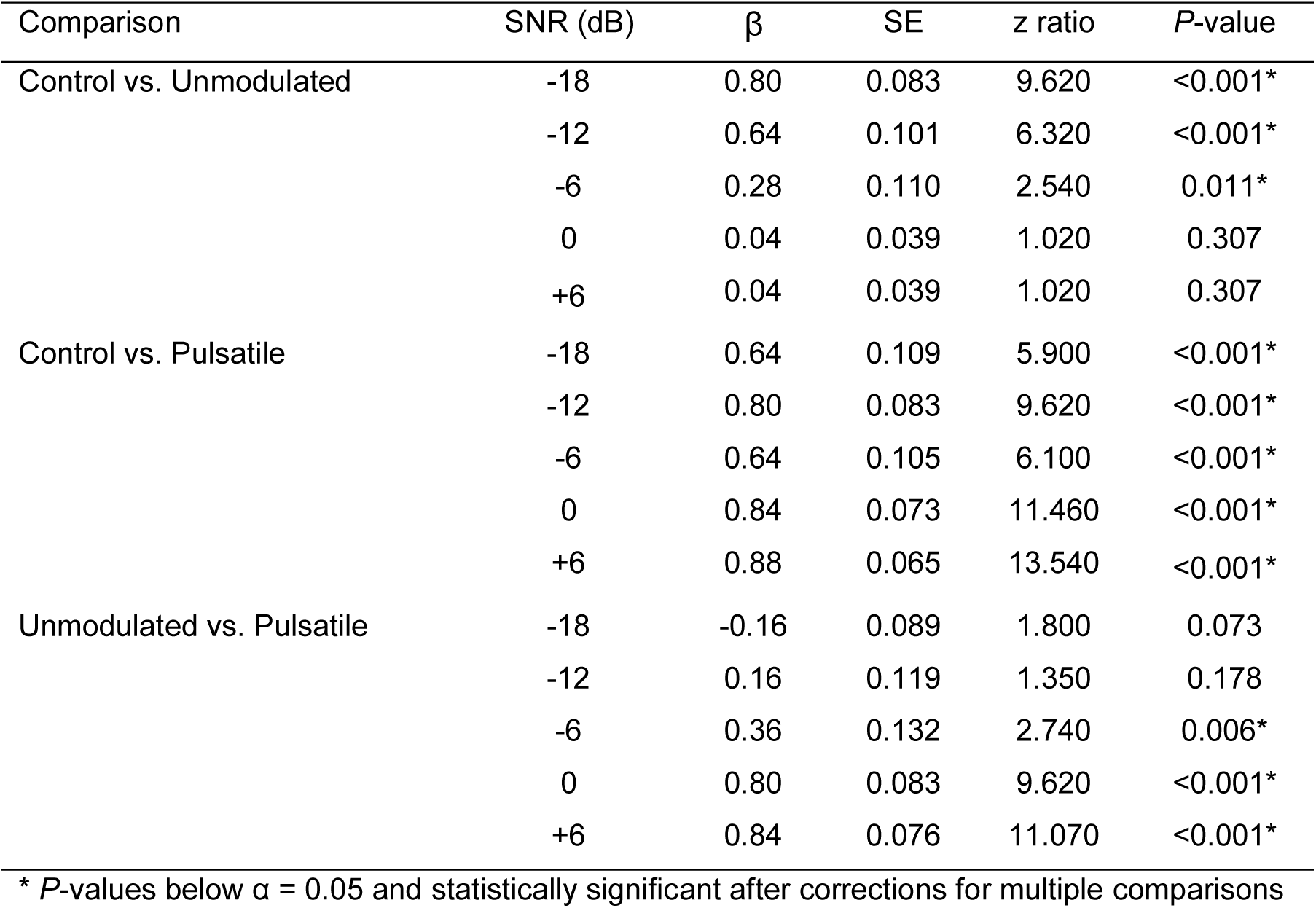
Outcomes of GEE pairwise comparisons between the three acoustic conditions (control, unmodulated, pulsatile) at each signal-to-noise ratio (SNR) from Experiment 1 (*n* = 50 per comparison). The table reports coefficients (β), standard errors (SE), z ratios, and associated *P*-values.

**Table S2.**
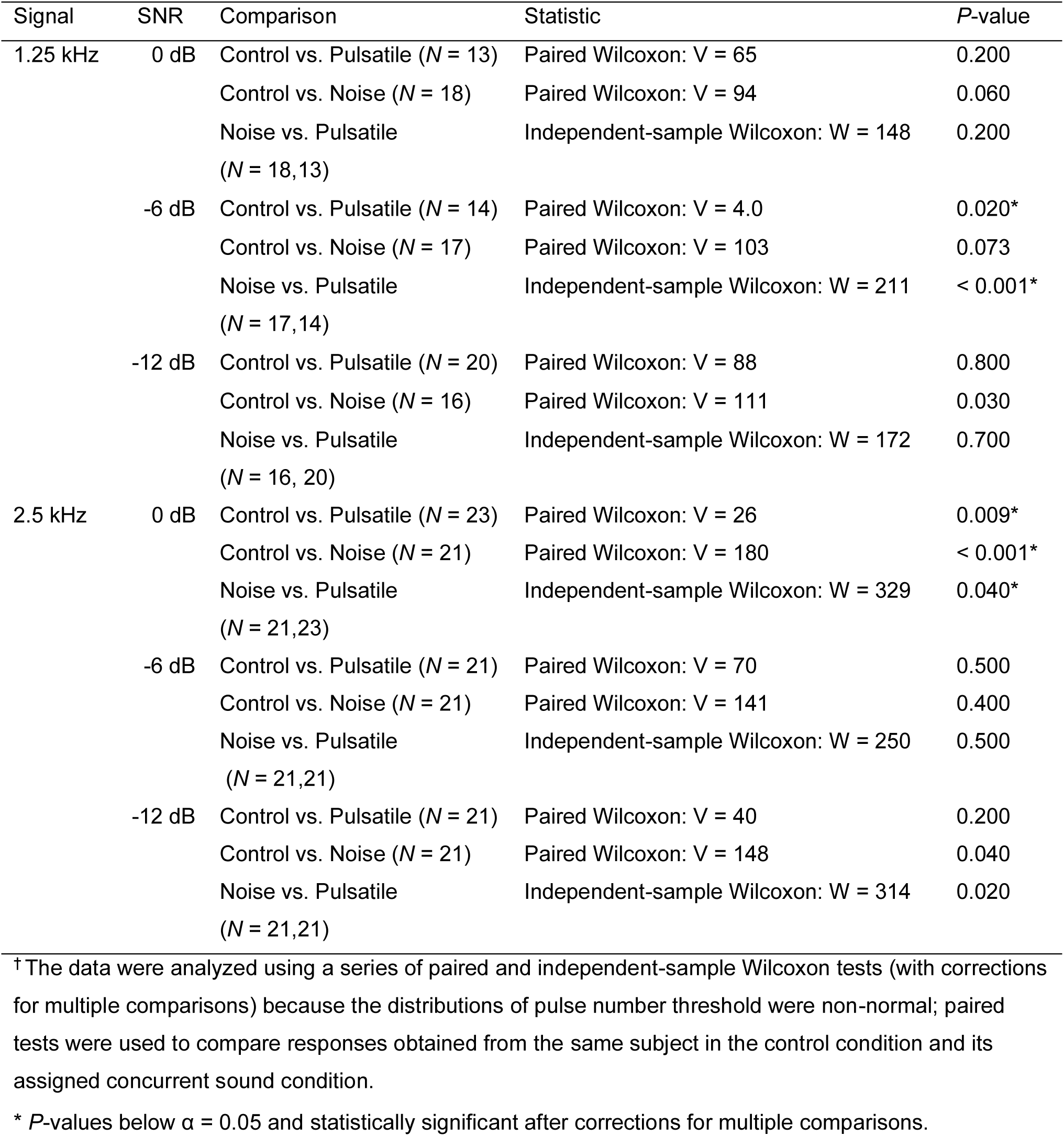
Outcomes of Wilcoxon signed rank tests^†^ comparing pulse number thresholds between the three acoustic conditions (control, pulsatile, and noise conditions) for the 1.25 kHz and 2.5 kHz signals in Experiment 2.

**Table S3.**
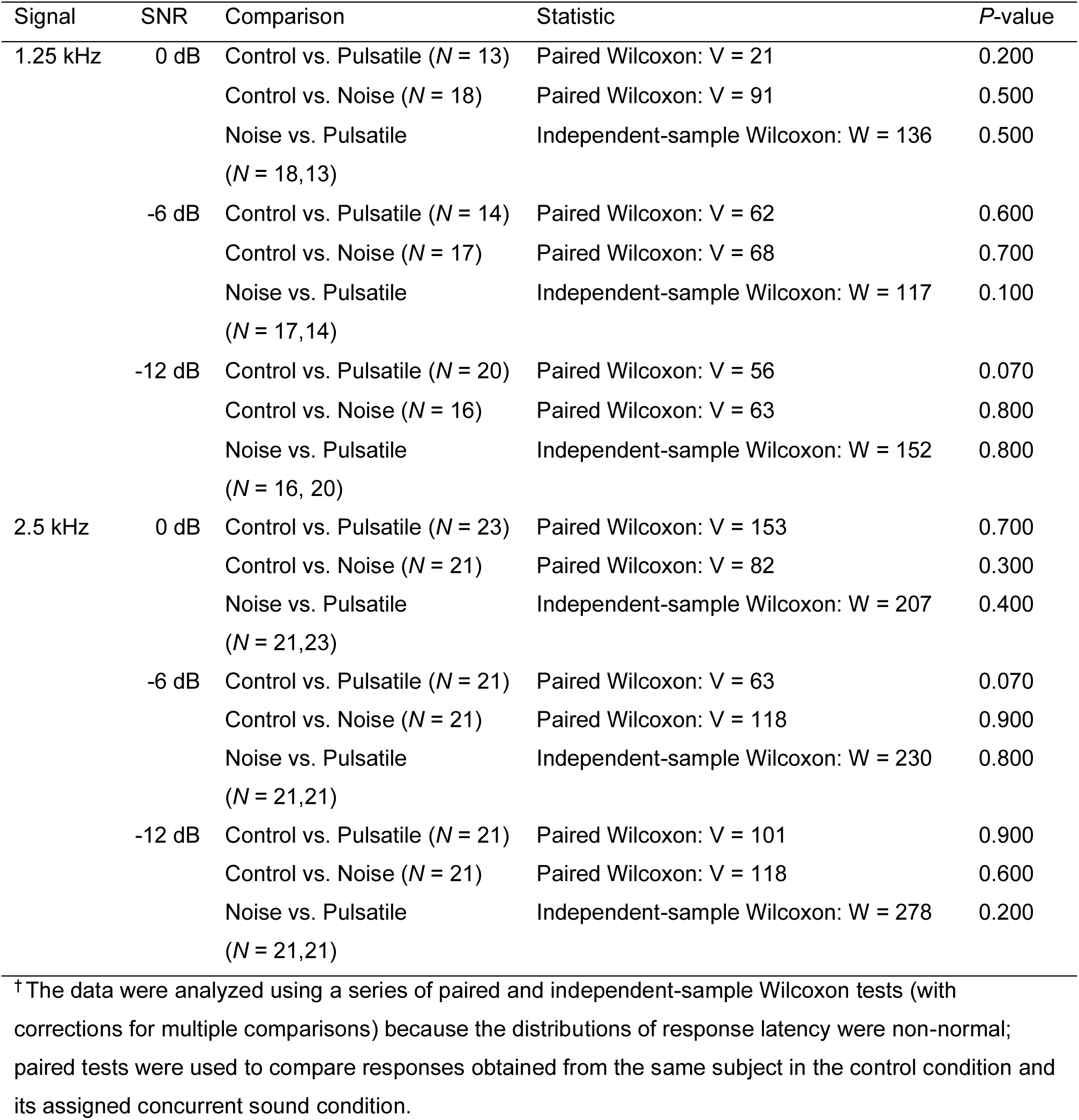
Outcomes of Wilcoxon signed rank tests^†^ comparing response latencies between the three acoustic conditions (control, pulsatile, and noise conditions) for the 1.25 kHz and 2.5 kHz signals in Experiment 2.

## Notes

### Competing Interest Statement

The authors have declared no competing interest.

### Summary of Updates

A new experiment (Experiment 1) has been added in this version

