## Supplemental Material for "Informational Masking Constrains Vocal Communication in Nonhuman Animals"

### Supplementary Materials for
### Informational Masking Constrains Vocal Communication in Nonhuman Animals

Gupta *et al.*

**This PDF file includes:**

Figures S1 to S4
Table S1 to S3

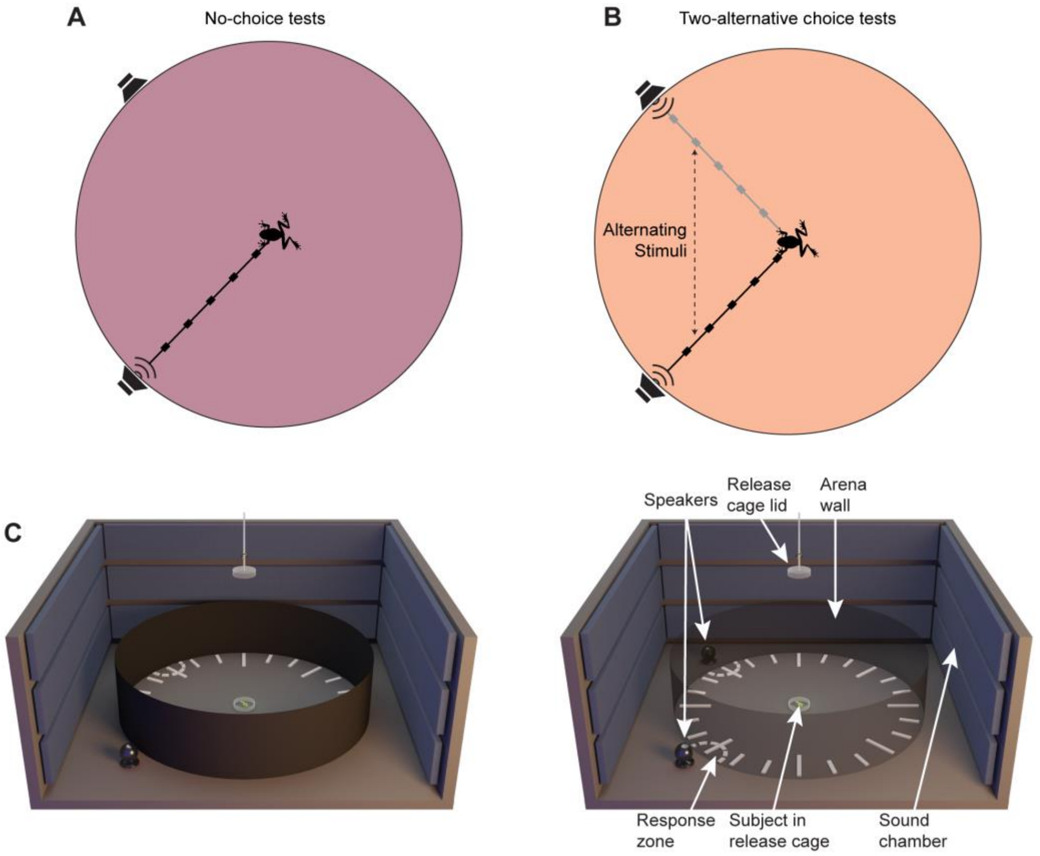


**Fig. S1. Schematic illustrations of testing protocols and experimental setup. (A,B)** Overhead schematics (not to scale) of the testing arena illustrating a single stimulus, no-choice test **(A)** and a two-alternative choice test **(B)**. **(A)** In no-choice tests, a single repeated stimulus was presented from one of the two speakers separated by 90º around the test arena. **(B)** In two alternative choice tests, two sequences of repeating stimuli (e.g., a signal alone or a signal plus concurrent sound) were presented from the two speakers; stimuli in the two sequences alternated in time such that there were equal intervals of silence preceding and following each stimulus presentation. To minimize any side bias, we randomized which speaker broadcast which stimulus across subjects. Alternating stimulus sequences are illustrated by gray and black schematic waveforms. **(C)** A 3D schematic of the circular test arena used to conduct phonotaxis tests with female treefrogs shown with visually opaque walls (Left) with walls rendered transparent (Right) to facilitate visualization of both playback speakers. The test arena (2-m diameter, 60-cm height) was located inside a semi anechoic chamber and was made from hardware cloth covered in black fabric. It was acoustically transparent but visually opaque. Acoustic stimuli were broadcast from one or two speakers positioned on the floor outside the arena wall and pointed toward a subject release cage at the center of the arena. The lid of the release cage could be remotely lifted from outside the chamber to allow the subject to respond by approaching a response zone. A response was scored when the subject entered a response zone. Responses were recorded using an infrared sensitive camera mounted above the center of the test arena (not shown) and viewed in real time by one or two experimenters on a monitor located outside the arena.

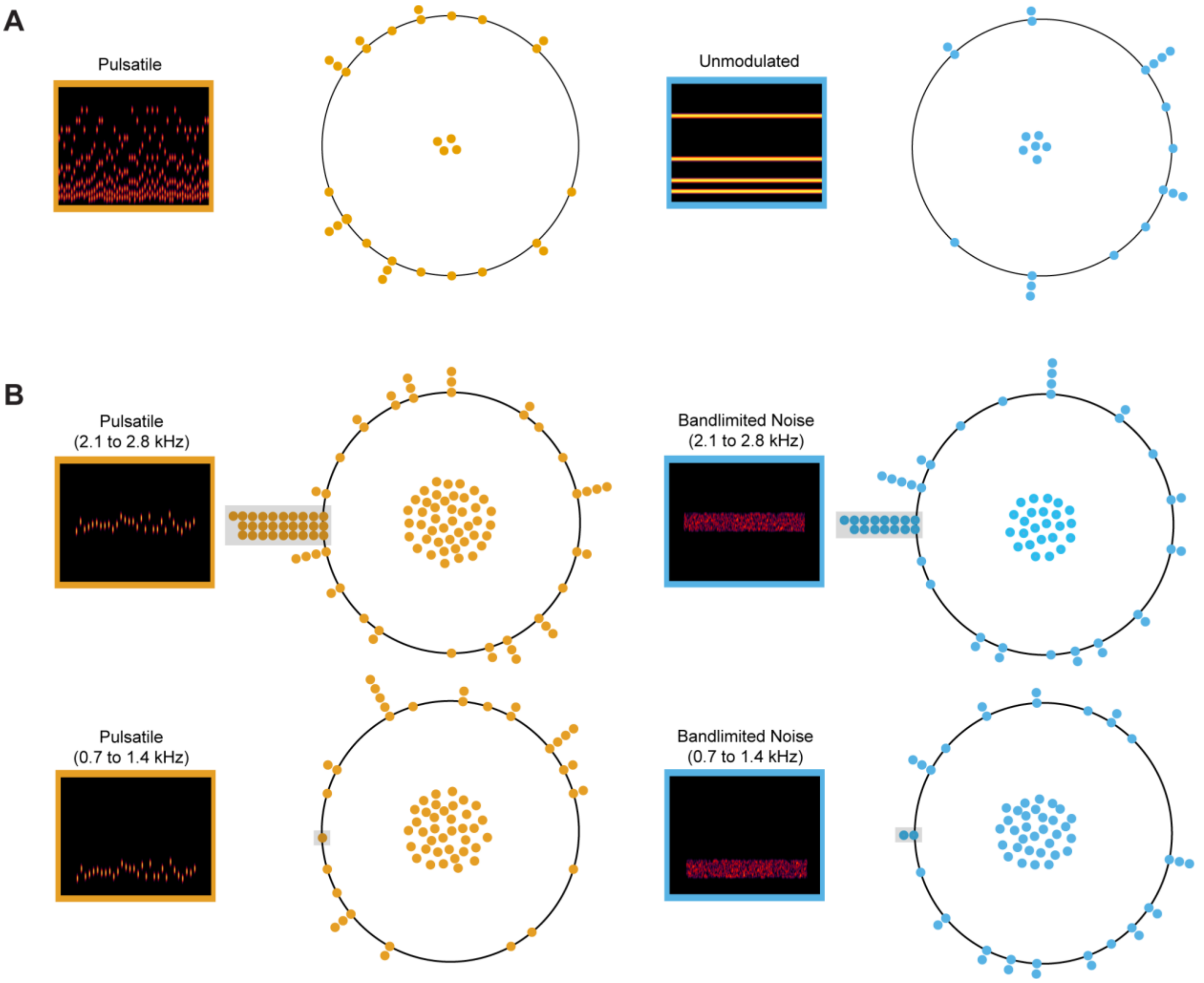


**Fig. S2. Concurrent sounds presented by themselves were not behaviorally aversive**. Points on each circle show where subjects tested in sham trials (i.e., no target signal) first touched the wall of the circular test arena relative to the playback speaker positioned at 9 o'clock around the circle. Individuals that never exited the release cage are depicted as points at the center of each circle and are excluded from the circular statistics (Rayleigh tests of uniformity) reported here. Insets show spectrograms illustrating the stimulus used in each sham trial. **(A)** The pulsatile and unmodulated sounds used in Experiment 1 were not behaviorally aversive, eliciting neither positive nor negative phonotaxis (pulsatile sound: r = 0.260, *P* = 0.172; unmodulated sound: r = 0.382, *P* = 0.061). **(B)** The pulsatile sound and bandlimited noise used in Experiments 2 and 3 were also not behaviorally aversive. Some individuals exhibited positive phonotaxis (gray shaded regions) to the pulsatile sound (*n* = 28 of 114) and bandlimited noise (*n* = 15 of 72) in a frequency range (2.1 to 2.8 kHz) transduced by the basilar papilla (BP; **B**, *top row*). After excluding these individuals, the remaining subjects exhibited neither positive nor negative phonotaxis (pulsatile sound: r = 0.021, *P* = 0.981; bandlimited noise: r = 0.068, *P* = 0.861). In contrast, few subjects exhibited positive phonotaxis to the pulsatile sound (*n* = 1 of 70) and bandlimited noise (*n* = 2 of 65) in a frequency range (0.7 to 1.4 kHz) transduced by the amphibian papilla (AP; **B,** *bottom row*). After excluding these individuals, the remaining subjects exhibited neither positive nor negative phonotaxis (pulsatile sound: r = 0.286, *P* = 0.071; bandlimited noise: r = 0.131, *P* = 0.590). All individuals that exhibited positive phonotaxis were excluded Experiment 2 to avoid introducing ambiguity into estimates of pulse number thresholds. A significantly greater proportion of subjects exhibited positive phonotaxis toward stimuli presented in the BP range than in the AP range (two proportion z-test: pulsatile, $\chi^2 = 15.8$, $P < 0.001$; band-limited noise, $\chi^2 = 8.5$, $P = 0.002$). This differential response to concurrent sounds that were similar in all aspects other than their frequency range corroborates previous findings showing that high-frequency sounds are perceptually more salient than low-frequency sounds in *H. chrysoscelis*.

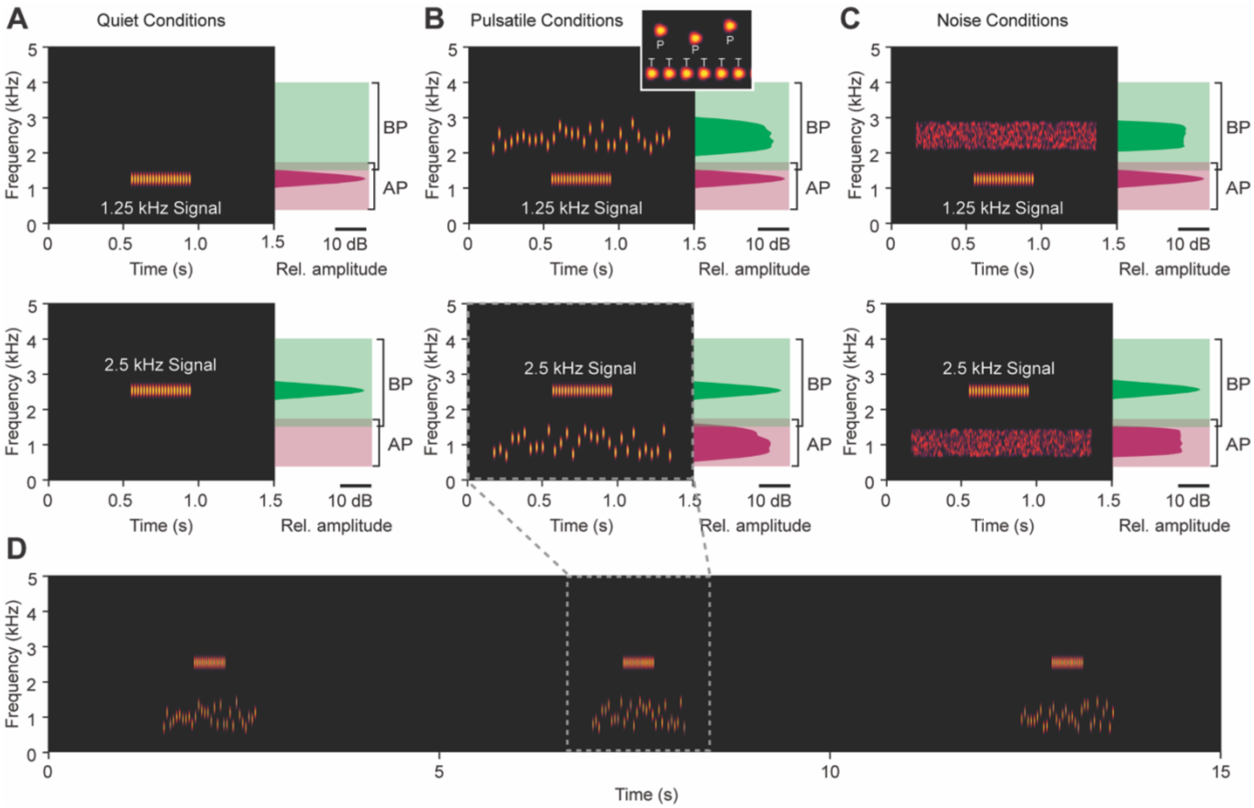


**Fig. S3. Examples of stimuli used to investigate call recognition in Experiments 2 and call discrimination in Experiment 3.** The 1.25 kHz and 2.5 kHz target signals (20 pulses) are shown separately in the (**A**) control, (**B**) pulsatile, and (**C**) noise conditions. Each plot in **A-C** shows a spectrogram (frequency × time) and power spectrum (frequency × relative amplitude) of one repetition of a target signal either in control (**A**) or with its associated concurrent sound (**B, C**). The inset in (**B**) illustrates how the pulses (P) of the concurrent sound in the pulsatile condition were temporally interleaved with every other pulse in the target signal (T). Power spectra are depicted relative to the approximate frequency ranges of the basilar papilla (BP; green) and amphibian papilla (AP; pink), the two sensory papillae in the frog inner ear that transduce airborne sound frequencies. Based on recordings of the auditory brainstem response in gray treefrogs, the lower spectral component (~1.25 kHz) is transduced primarily by the AP (range of sensitivity ~0.4 to 1.75 kHz), and the higher component (~2.5 kHz) is transduced primarily by the BP (range of sensitivity ~1.5 to 4.0 kHz). As illustrated, there is minimal overlap in the tuning of the two sensory papillae. By specifying the carrier frequency of the target signal at either 1.25 kHz or 2.5 kHz and limiting concurrent sounds to the opposite frequency range, the stimuli were designed such that target signals and concurrent sounds would primarily stimulate different inner ear papillae. The concurrent sound in the pulsatile condition was a series of random-frequency pulses (25 pulses/s) that were interleaved between every other pulse in a target signal (50 pulses/s). The narrowband noise in the noise condition had the same spectral content and long-term sound pressure level as the pulsatile concurrent sound. In both conditions with concurrent sounds, target signals were temporally centered within the concurrent sound, which was gated on and off around the signal. (**D**) A 15-s segment of a stimulus using in Experiments 1 and 2. During phonotaxis tests, stimuli were repeated at a rate of 11 calls/min for up to 5 minutes or until a subject responded.

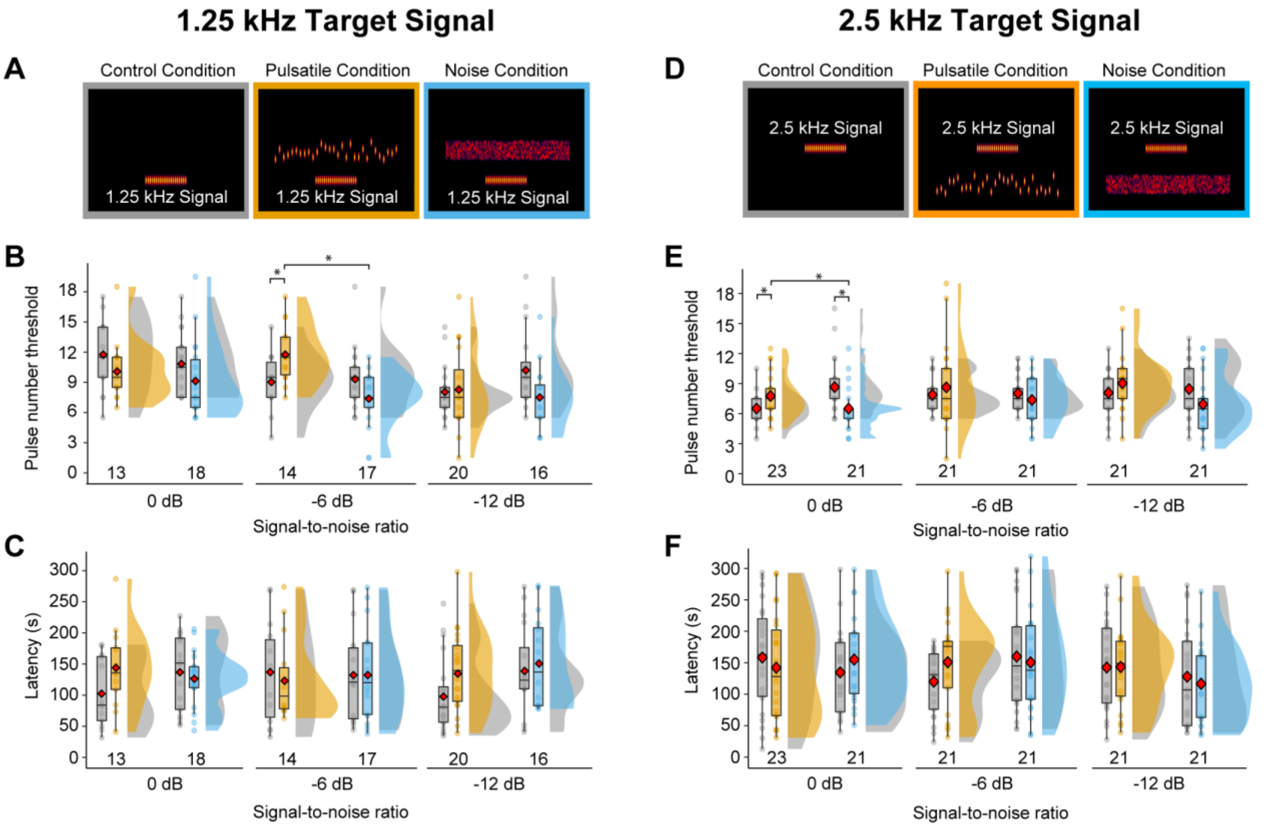


**Fig. S4. Experiment 2: Temporally modulated concurrent sounds had limited impact on pulse number thresholds and response latencies.** (**A**) Spectrograms illustrating the 1.25 kHz signal with 20 pulses in the control (*gray*), pulsatile (*orange*), and noise (*blue*) conditions. (**B**) Pulse number thresholds for responding to 1.25 kHz signals measured using an adaptive tracking procedure and defined as the minimum number of pulses required to elicit positive phonotaxis. For each subject tested at each SNR, separate thresholds were determined within subjects in the control condition and in either the pulsatile condition or the noise condition. Box plots depict the first, second (median), and third quartiles, and the mean (red diamond) values; whiskers depict 1.5 times the interquartile range and points depict individual data points. The two overlapping distributions are kernel density representations of the data that are depicted in the corresponding box plots. For each SNR, the two box plots and distributions on the left show data for the control and pulsatile conditions that were paired within subjects, whereas the two box plots and distributions on the right show data for the control and noise conditions that were paired within a different group of subjects. Sample sizes are indicated at the base of each dataset. At a SNR of -6 dB, pulse number thresholds were significantly higher (*$P < 0.05$) in the pulsatile condition compared with the paired control condition and the noise condition (Table S1). (**C**) Response latencies for the 1.25 kHz signal, computed as the time to respond to the target signal having the lowest number of pulses that elicited positive phonotaxis. There were no significant differences in response latencies. Box plots and distributions as in (**B**). (**D**) Spectrograms illustrating the 2.5 kHz signal with 20 pulses in the control (*gray*), pulsatile (*orange*), and noise (*blue*) conditions. (**E**) Pulse number thresholds for responding to 2.5 kHz signals. At a SNR of 0 dB, pulse number thresholds were significantly higher in the pulsatile condition compared with both the paired control condition and the noise condition; they were significantly lower in the noise condition compared with the paired control condition (Table S2). (**F**) Response latencies for 2.5 kHz signal. There were no significant differences in response latencies (Table S3).

**Table S1.** Outcomes of GEE pairwise comparisons between the three acoustic conditions (control, unmodulated, pulsatile) at each signal-to-noise ratio (SNR) from Experiment 1 (*n* = 50 per comparison). The table reports coefficients (β), standard errors (SE), z ratios, and associated *P*-values.

| Comparison | SNR (dB) | β | SE | z ratio | *P*-value |
|---|---|---|---|---|---|
| Control vs. Unmodulated | -18 | 0.80 | 0.083 | 9.620 | <0.001* |
| | -12 | 0.64 | 0.101 | 6.320 | <0.001* |
| | -6 | 0.28 | 0.110 | 2.540 | 0.011* |
| | 0 | 0.04 | 0.039 | 1.020 | 0.307 |
| | +6 | 0.04 | 0.039 | 1.020 | 0.307 |
| Control vs. Pulsatile | -18 | 0.64 | 0.109 | 5.900 | <0.001* |
| | -12 | 0.80 | 0.083 | 9.620 | <0.001* |
| | -6 | 0.64 | 0.105 | 6.100 | <0.001* |
| | 0 | 0.84 | 0.073 | 11.460 | <0.001* |
| | +6 | 0.88 | 0.065 | 13.540 | <0.001* |
| Unmodulated vs. Pulsatile | -18 | -0.16 | 0.089 | 1.800 | 0.073 |
| | -12 | 0.16 | 0.119 | 1.350 | 0.178 |
| | -6 | 0.36 | 0.132 | 2.740 | 0.006* |
| | 0 | 0.80 | 0.083 | 9.620 | <0.001* |
| | +6 | 0.84 | 0.076 | 11.070 | <0.001* |

* *P*-values below α = 0.05 and statistically significant after corrections for multiple comparisons

**Table S2.** Outcomes of Wilcoxon signed rank tests† comparing pulse number thresholds between the three acoustic conditions (control, pulsatile, and noise conditions) for the 1.25 kHz and 2.5 kHz signals in Experiment 2.

| Signal | SNR | Comparison | Statistic | *P*-value |
|---|---|---|---|---|
| 1.25 kHz | 0 dB | Control vs. Pulsatile (*N* = 13) | Paired Wilcoxon: V = 65 | 0.200 |
| | | Control vs. Noise (*N* = 18) | Paired Wilcoxon: V = 94 | 0.060 |
| | | Noise vs. Pulsatile (*N* = 18,13) | Independent-sample Wilcoxon: W = 148 | 0.200 |
| | -6 dB | Control vs. Pulsatile (*N* = 14) | Paired Wilcoxon: V = 4.0 | 0.020* |
| | | Control vs. Noise (*N* = 17) | Paired Wilcoxon: V = 103 | 0.073 |
| | | Noise vs. Pulsatile (*N* = 17,14) | Independent-sample Wilcoxon: W = 211 | < 0.001* |
| | -12 dB | Control vs. Pulsatile (*N* = 20) | Paired Wilcoxon: V = 88 | 0.800 |
| | | Control vs. Noise (*N* = 16) | Paired Wilcoxon: V = 111 | 0.030 |
| | | Noise vs. Pulsatile (*N* = 16, 20) | Independent-sample Wilcoxon: W = 172 | 0.700 |
| 2.5 kHz | 0 dB | Control vs. Pulsatile (*N* = 23) | Paired Wilcoxon: V = 26 | 0.009* |
| | | Control vs. Noise (*N* = 21) | Paired Wilcoxon: V = 180 | < 0.001* |
| | | Noise vs. Pulsatile (*N* = 21,23) | Independent-sample Wilcoxon: W = 329 | 0.040* |
| | -6 dB | Control vs. Pulsatile (*N* = 21) | Paired Wilcoxon: V = 70 | 0.500 |
| | | Control vs. Noise (*N* = 21) | Paired Wilcoxon: V = 141 | 0.400 |
| | | Noise vs. Pulsatile (*N* = 21,21) | Independent-sample Wilcoxon: W = 250 | 0.500 |
| | -12 dB | Control vs. Pulsatile (*N* = 21) | Paired Wilcoxon: V = 40 | 0.200 |
| | | Control vs. Noise (*N* = 21) | Paired Wilcoxon: V = 148 | 0.040 |
| | | Noise vs. Pulsatile (*N* = 21,21) | Independent-sample Wilcoxon: W = 314 | 0.020 |

† The data were analyzed using a series of paired and independent-sample Wilcoxon tests (with corrections for multiple comparisons) because the distributions of pulse number threshold were non-normal; paired tests were used to compare responses obtained from the same subject in the control condition and its assigned concurrent sound condition.

* *P*-values below α = 0.05 and statistically significant after corrections for multiple comparisons.

**Table S3.** Outcomes of Wilcoxon signed rank tests† comparing response latencies between the three acoustic conditions (control, pulsatile, and noise conditions) for the 1.25 kHz and 2.5 kHz signals in Experiment 2.

| Signal | SNR | Comparison | Statistic | *P*-value |
|---|---|---|---|---|
| 1.25 kHz | 0 dB | Control vs. Pulsatile (*N* = 13) | Paired Wilcoxon: V = 21 | 0.200 |
| | | Control vs. Noise (*N* = 18) | Paired Wilcoxon: V = 91 | 0.500 |
| | | Noise vs. Pulsatile (*N* = 18,13) | Independent-sample Wilcoxon: W = 136 | 0.500 |
| | -6 dB | Control vs. Pulsatile (*N* = 14) | Paired Wilcoxon: V = 62 | 0.600 |
| | | Control vs. Noise (*N* = 17) | Paired Wilcoxon: V = 68 | 0.700 |
| | | Noise vs. Pulsatile (*N* = 17,14) | Independent-sample Wilcoxon: W = 117 | 0.100 |
| | -12 dB | Control vs. Pulsatile (*N* = 20) | Paired Wilcoxon: V = 56 | 0.070 |
| | | Control vs. Noise (*N* = 16) | Paired Wilcoxon: V = 63 | 0.800 |
| | | Noise vs. Pulsatile (*N* = 16, 20) | Independent-sample Wilcoxon: W = 152 | 0.800 |
| 2.5 kHz | 0 dB | Control vs. Pulsatile (*N* = 23) | Paired Wilcoxon: V = 153 | 0.700 |
| | | Control vs. Noise (*N* = 21) | Paired Wilcoxon: V = 82 | 0.300 |
| | | Noise vs. Pulsatile (*N* = 21,23) | Independent-sample Wilcoxon: W = 207 | 0.400 |
| | -6 dB | Control vs. Pulsatile (*N* = 21) | Paired Wilcoxon: V = 63 | 0.070 |
| | | Control vs. Noise (*N* = 21) | Paired Wilcoxon: V = 118 | 0.900 |
| | | Noise vs. Pulsatile (*N* = 21,21) | Independent-sample Wilcoxon: W = 230 | 0.800 |
| | -12 dB | Control vs. Pulsatile (*N* = 21) | Paired Wilcoxon: V = 101 | 0.900 |
| | | Control vs. Noise (*N* = 21) | Paired Wilcoxon: V = 118 | 0.600 |
| | | Noise vs. Pulsatile (*N* = 21,21) | Independent-sample Wilcoxon: W = 278 | 0.200 |

† The data were analyzed using a series of paired and independent-sample Wilcoxon tests (with corrections for multiple comparisons) because the distributions of response latency were non-normal; paired tests were used to compare responses obtained from the same subject in the control condition and its assigned concurrent sound condition.
